# Single-cell omics reveal distinct gene regulatory dynamics underpinning embryonic and extraembryonic lineage functions during pig blastocyst development

**DOI:** 10.1101/2025.06.15.657618

**Authors:** Adrien Dufour, Marie-Noëlle Rossignol, Patrick Manceau, Yoann Bailly, Stéphane Ferchaud, Marie-José Mercat, Ali G Turhan, Sarah Djebali, Sylvain Foissac, Jérôme Artus, Hervé Acloque

## Abstract

The late-stage development of the blastocyst before implantation is a unique feature of ungulates. During this period, the epiblast proliferates and remains pluripotent for several days before gastrulation begins. Simultaneously, extra-embryonic tissues undergo significant growth, elongating to several tens of centimeters. However, the mechanisms that regulate and synchronize these processes remain poorly understood. In this study, we analyzed transcriptomic and epigenomic data at the single-cell level from early and late porcine blastocyst cells, spanning from the hatched blastocyst stage (E7) to early (E9) and late ovoid blastocyst (E11). We characterized 15,370 blastocyst cells, clustering them into distinct embryonic and extra-embryonic populations with specific chromatin accessibility landscapes. For each population, we inferred gene regulatory networks based on enhancer-driven gene regulation modules (eRegulons) and validated them through motif occupancy visualization. Our findings reveal strong dynamics in gene regulatory module activity within extra-embryonic tissues at the onset of blastocyst elongation, while gene regulatory activity in the epiblast remains relatively stable.

## Introduction

Pigs are one of the main sources of animal products for humans and the most widely consumed meat worldwide. They also serve as an alternative to rodent models for studying early embryo development^1^ and as an attractive biomedical model for human pathology^2^. Pig early embryonic development follows a pattern similar to other mammals. After fertilization, the embryo undergoes several cleavage divisions, followed by compaction and the formation of the blastocyst, composed of an inner cell mass (ICM) and an outer trophectoderm (TE). By 6-day post-fertilization (dpf), the ICM differentiates into the hypoblast or primitive endoderm (HYPO) and the epiblast (EPI). Unlike in rodents or primates, the porcine blastocyst begins to elongate from 7-dpf, transitioning through ovoid, tubular, and filamentous stages. During this process, the trophectoderm covering the epiblast (EPI), known as Rauber’s layer, disappears.

We and others have explored the development of the pig pre-implantation embryo at the transcriptomics single-cell (scRNA-seq) level, before and during gastrulation^3–6^. These studies have highlighted the role of IL6-JAK-STAT signaling in ICM and TE formation. They also have shown that the transition of pluripotency status from a naive-like to a prime-like state is associated with changes in active signaling pathways and pluripotency marker expression. Indeed, naive-like pluripotency at embryonic-day-5 (E5) is characterised by high activity of IL6-STAT3 and PI3K-AKT signaling pathways associated with the expression of naive pluripotency markers (*KLF4*, *ESRRB*, *STAT3*). Conversely, from E7 onwards, pluripotency shifts to a primed-like state marked by increased activity of TGFβ-SMAD2/3 signaling pathway associated with the expression of primed pluripotency genes (*NANOG*, *DNMT3B*, *OTX*2). This transition is also accompanied by epigenetic changes through DNA and histone modifications, including X chromosome inactivation in female EPI cells, and a metabolic switch from OXPHOS to glycolysis^3–5^.

Interestingly, our previous single-cell study identified new subpopulations within extra-embryonic cell lineages. These include cells that secrete molecules required for implantation, such as interleukin-1 beta (*IL1B*), and a putative trophoblast stem cell population expressing *LRP2*, which later contributes to the development of the embryonic placenta^5^. Our data also confirmed that extra-embryonic subpopulations play key roles in the metabolism of fatty acids, vitamins, nutrient transport and conceptus elongation^5^. By integrating this dataset, we identified several transcription factors (TFs) and their associated regulons^7^, as well as ligand-receptor pairs related to these populations. This integration allowed us to predict the upstream signalling pathways governing gene regulatory modules.

While these studies provide a better understanding of early pig embryonic development, they have not explored the epigenetic changes occurring during these stages. Such modifications are thought to be critical during this developmental window, as they likely regulate gene expression dynamics, influence lineage commitment, and establish the epigenetic landscape necessary for proper embryo implantation and subsequent development. Modifications like *de novo* DNA methylation in the epiblast, are suggested by the increased expression of genes such as *DNTM3B*, *DNMT3A* and *HELLS* in early pig blastocysts^4^. These questions have been addressed in other mammals, such as mice, using bulk ATAC-seq, histone profiling and scATAC-seq^8^. More recently, single-cell multiomics (combining RNA-seq and ATAC-seq in the same cells) has emerged as a powerful approach for investigating gene regulatory networks (GRN), with new analytical methods such as SCENIC+ recently applied to the fly brain^9^, though they remain underutilized in mammalian embryos.

Here, we produced and analysed a single-cell multiomics dataset of pig preimplantation embryos across key developmental stages, from hatched blastocyst (E7) to spheroid/early ovoid blastocyst (E9) and late ovoid blastocyst (E11) (Figure 1A). We characterised 15,370 embryonic cells, which we clustered into distinct embryonic and extra-embryonic cell populations associated with specific chromatin accessibility landscapes. For each cell population, we then inferred GRNs based on enhancer-driven gene regulation modules (eRegulons)^10^ and validated them through motif occupancy visualisation. We detected known and new eRegulons for each cell population, and observed a strong dynamics in eRegulons activity within extra-embryonic tissues (TE and HYPO) at the onset of blastocyst elongation, while the activity of gene regulatory modules remained relatively constant in the epiblast.

**Figure 1.**
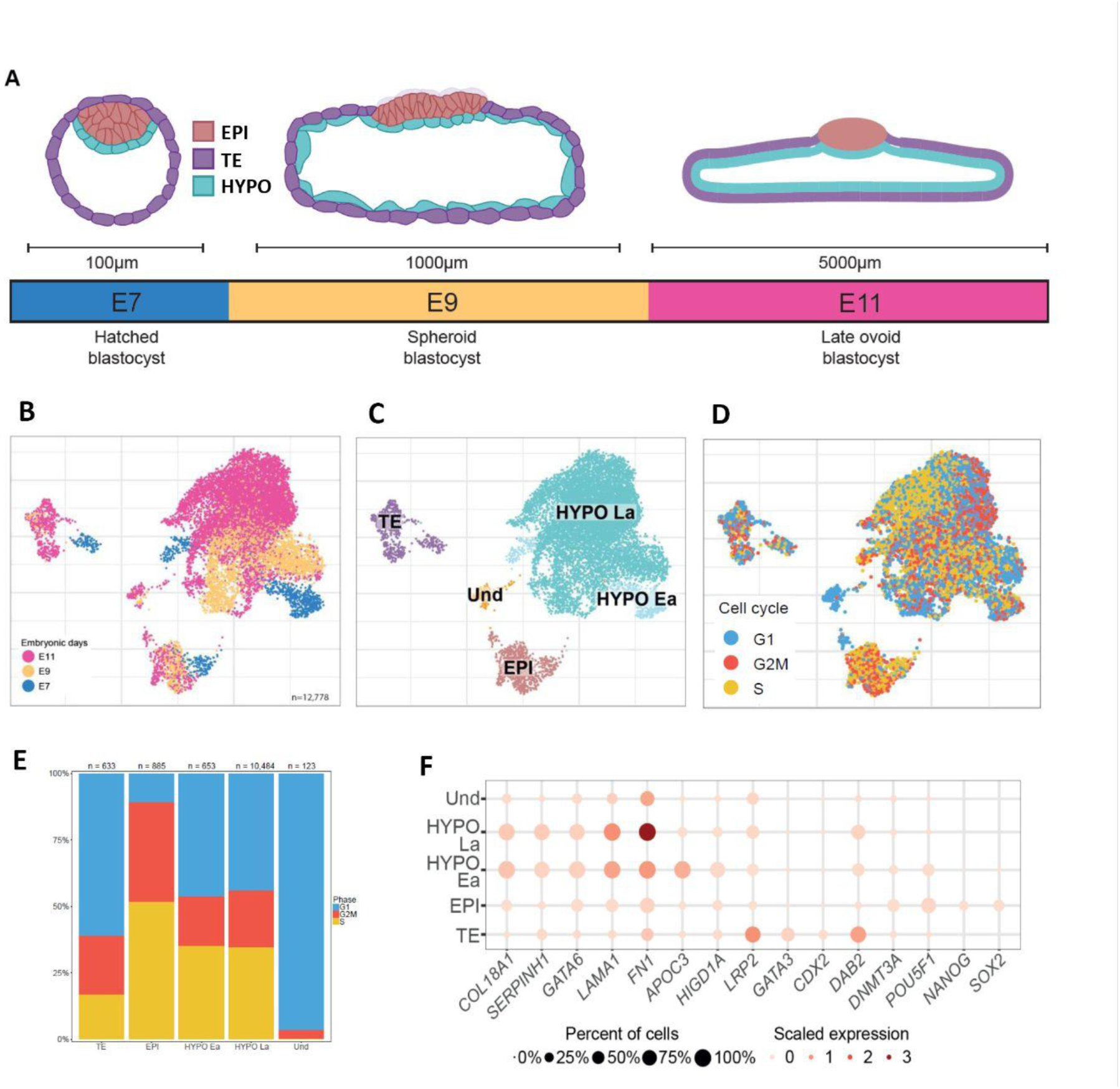
Single-cell multiomics sequencing of pig pre-implantation embryos. **A**. Schematic view of pig embryo morphology for embryonic day (E) E7, E9 and E11. Cells from the EPI, TE and HYPO are represented respectively in red, purple and light blue. **B**. UMAP representation of all single-cells passing quality control coloured by embryonic stage (E7 in blue, E9 in orange and E11 in purple). **C**. UMAP representation of the assigned population : EPI (red), HYPO Late (turquoise), HYPO Early (light blue), TE (purple) and Und (undetermined, orange). **D.** UMAP representation of cell cycle assignment of each cell with respect to the three different phases of the cell cycle: Gap1 (G1, in blue), Synthesis (S, in yellow), Gap2 and mitosis (G2/M, in red). Cells have been assigned to the cell cycle phase obtaining the best score. **E.** Proportion of cells in G1 (blue), G2/M (red) and S (yellow) in each cluster. **F**. Dot plot of gene expression in cell clusters for selected markers. Red colour gradient represents the relative expression level across clusters and dot size represents the percentage of cells in that cluster expressing the gene.

## Results

### Joint transcriptomic and epigenomic profiling of embryonic and extraembryonic lineages during pig blastocyst development

We generated a paired transcriptomic and chromatin accessibility profile using the 10x Multiome technology from single nuclei of pig preimplantation embryos collected at E7, E9 and E11 (Figure 1A, Supplementary Table 1). A total of 12,778 cells passed quality control for both data modalities (Figure 1B, Supplementary Figure 1, Supplementary Table 2). We identified four cell clusters that were classified as TE, HYPO early/late and EPI according to the expression of specific lineage markers (Figure 1C). To confirm our assignment, we plotted, on the same UMAP, another assignment based on population signatures obtained from other scRNA-seq datasets obtained from pig preimplantation embryos^5,6^ (Supplementary Figure 2). Within the hypoblast, we identified two clusters: early hypoblast (HYPO Ea) and late hypoblast (HYPO La). HYPO Ea cells express markers such as *APOC3* and *HIGD1A* (Figure 1F), which we previously associated to early and intermediate hypoblast (HYPO Ea and HYPO In populations, Supplementary Figure 2C)^5^ and are exclusively present at the earliest E7 and E9 stages. HYPO La cells express higher levels of *FN1* (Figure 1F), similarly to more mature hypoblast (HYPO Ma) cell populations (Supplementary Figure 2B-C)^5^. By performing cell cycle assignment using CellCycleScoring Seurat function (see Methods), we observed that Hypo Ea and Hypo La populations exhibit similar cell cycle profiles, with approximately 50% of cells in proliferative phases (G2/M and S) and the remaining 50% in G1 (Figure 1D-E). In this dataset, our clustering analysis identified a single trophectoderm (TE) cluster characterized by the expression of *LRP2*, *GATA3*, *CDX2*, and *DAB2* (Figure 1F). However, our cell assignment analysis clearly reveals the presence of distinct subpopulations previously described as TE LRP2 (TE Lr) and TE Mature2 (TE Ma2) (Supplementary Figure 2B-C)^5^. TE cells are proliferating less than the other populations with less than 40% of cells in proliferative phases (G2/M and S) (Figure 1D-E). EPI cells correspond to the Epiblast signature (yellow, Supplementary Figure 2B-C) and are associated with the expression of known pluripotency markers such as *NANOG*, *POU5F1*, *SOX2* and *DNMT3A* (Figure 1F). EPI cells display a strong proliferative profile, with over 80% of cells in G2/M and S phases (Figure 1D-E), confirming our previous observation^5^. One small population was named undetermined (Und) and does not express any markers of embryonic or extra-embryonic tissues. This population shows evidence of cell cycle exit, with most cells accumulating in G1 phase. Strikingly, the proportions of cells within the different populations (8% EPI, 87% HYPO, 5% TE) differ from our previous and other scRNA-seq studies on pig preimplantation embryos. This discrepancy likely originates from the nuclei isolation procedure^11^ while previous scRNA-seq studies typically analyze whole cells rather than isolated nuclei. Some cell types or subpopulations may be more fragile or susceptible to loss during the nuclei isolation process which can result in differences in observed cell proportions.

Each cell population exhibited distinct epigenetic landscapes. In the whole dataset, we identified 108,810 chromatin accessibility peaks (Supplementary File 1), located either within genes (intronic and exonic), promoter regions (500bp upstream and 100bp downstream) or distal regions (Figure 2A). 44% of these peaks are common to all populations while 15% are unique to HYPO and TE and 5% in EPI (Figure 2B). This observation prompted us to investigate to what extent the embryonic epigenetic landscape differs from that of differentiated somatic cells. By comparing our profiles of chromatin accessibility peaks with those from other somatic tissues^12^, we observed that the majority of these peaks are shared across cell types, with only a small proportion being specific to blastocyst tissues (Supplementary Figure 3 and Supplementary Table 3).

**Figure 2.**
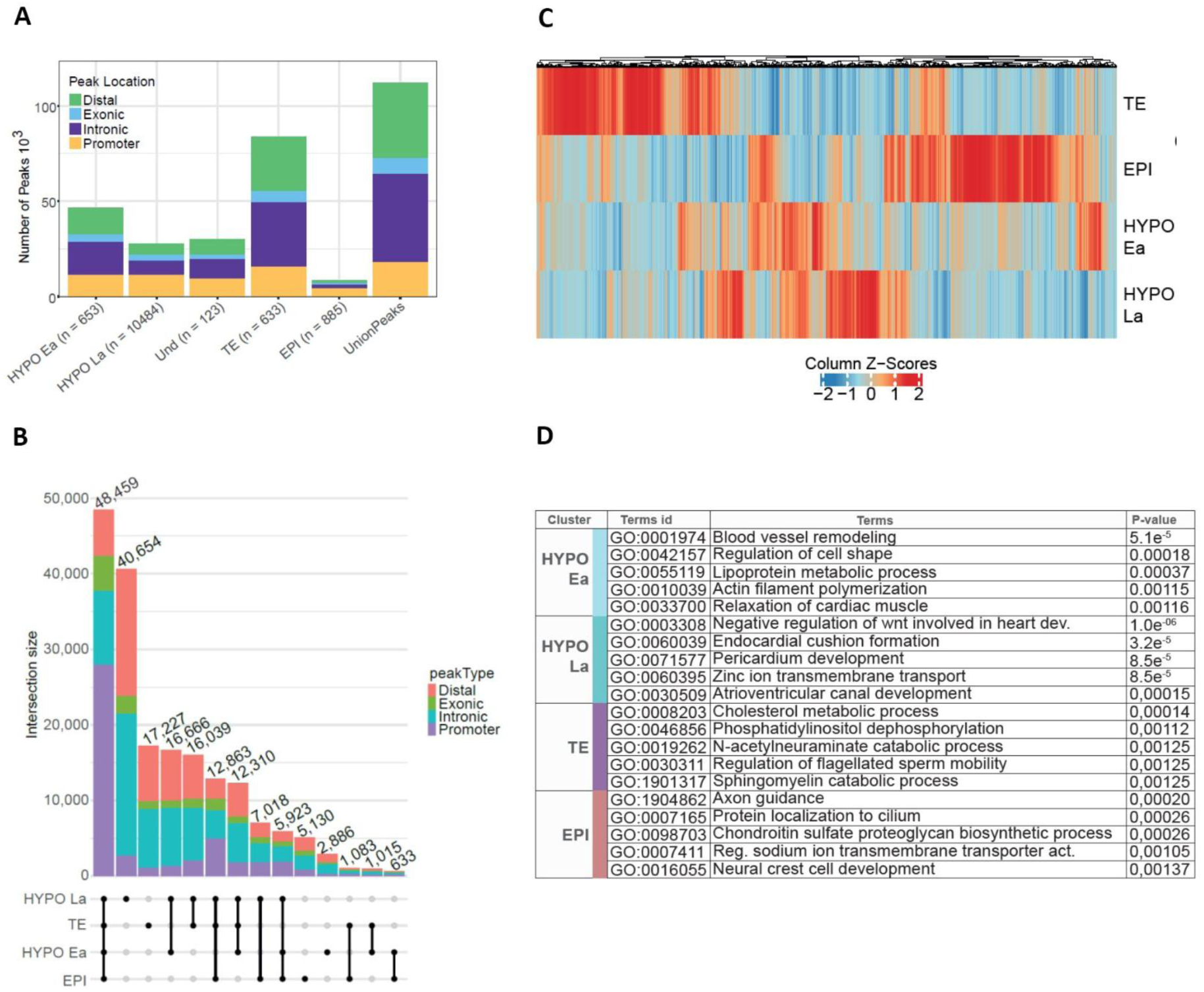
Epigenomic landscape of pig pre-implantation embryos. **A**. Barplot of peak quantification according to genome location in each cluster and resulting union of peaks. bp downstream from the promoter). **B.** Upset plot comparing the number of peaks overlap between cell populations. **C.** Heatmap reflecting differential promoter accessibility in each cell cluster (500bp upstream and 100bp downstream from the promoter). **D.** Gene Ontology: Biological Process enrichment in the significantly enriched DEGs in each cluster. The top 5 are shown.

We then determined whether blastocyst-specific regions reflect unique regulatory features associated with pluripotency and early lineage commitment. Using these peaks, we identified 14,016 differentially accessible peaks (DAPs) between the cell clusters including 980 located on promoter regions, highlighting specific epigenomic landscapes of each cell population (Figure 2B-C, Supplementary Table 4).

By integrating differentially expressed genes (DEGs) with differentially accessible promoter regions (DAPRs) across cell populations, we identified a strong correlation between chromatin accessibility profiles and the expression of established lineage markers (Figure 3, Supplementary Tables 5 and 6, Supplementary Figure 4). This analysis also provides functional insight into cis-regulatory elements active in pig blastocyst cells. Notably, regions of open chromatin were associated with elevated expression of key genes such as *APOC3*, *FN1*, *HIGD1A*, and *LAMA1* in the hypoblast (HYPO); *DAB2*, *GATA3*, and *CDX2* in the trophectoderm (TE); and *SOX2*, *NANOG*, and *POU5F1* in the epiblast (EPI) (Figure 3, Supplementary Figure 3).

**Figure 3.**
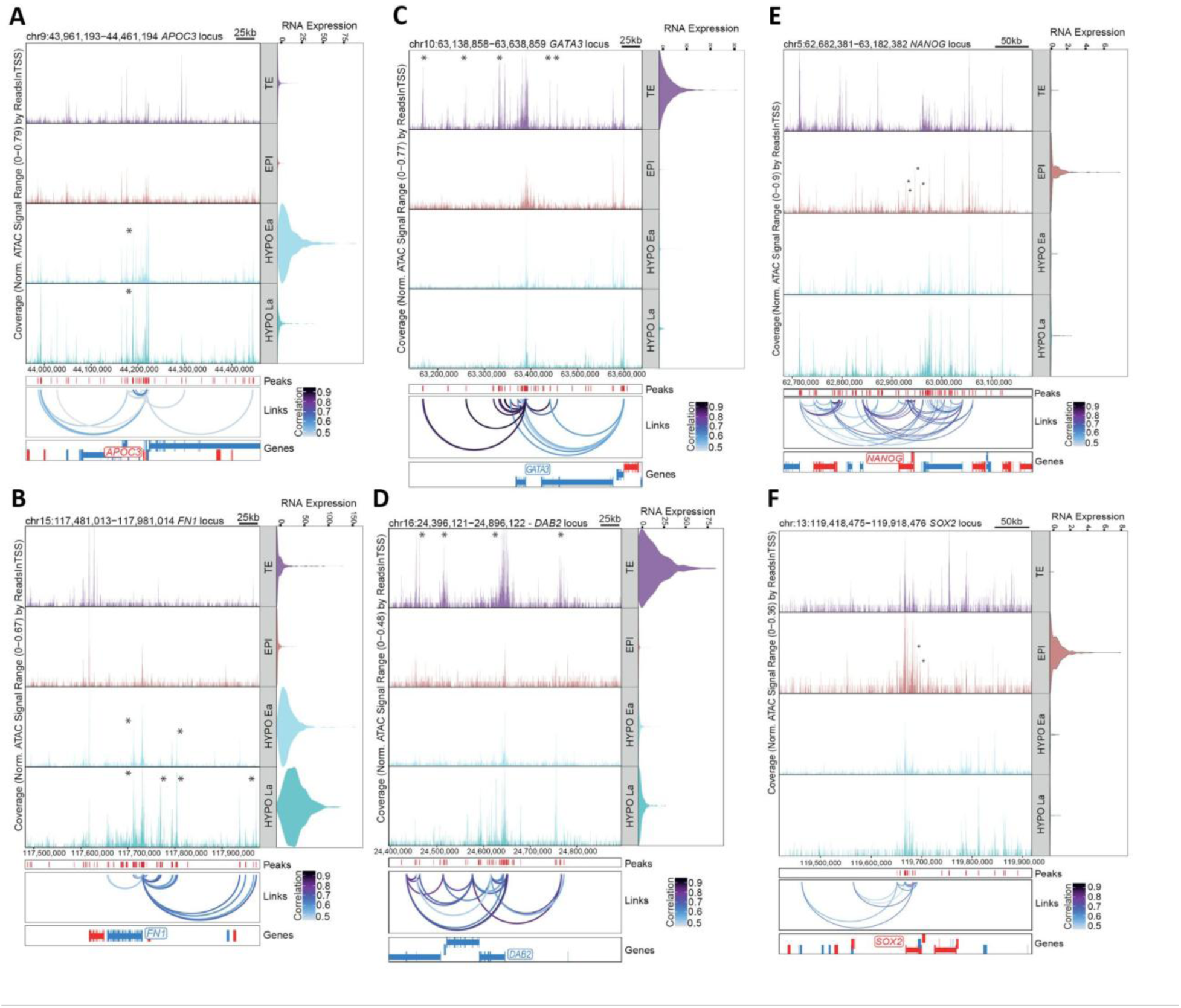
Profiles of peak-to-gene linkages for HYPO, TE and EPI markers in pig embryos. **A-F**. Coverage plots showing aggregated single-cell ATAC signals around *APOC3* (A), *FN1* (B), *GATA3* (C), *DAB2* (D), *NANOG* (E) and *SOX2* (F) loci within the different cell types. On the right of each coverage plot, violin plots show their expression levels for each cell type. Below each coverage plot, loops predicted by ArchR indicate peak-to-gene linkages identified on the full dataset (coloured by importance score). Stars indicate peak-to-gene links with a correlation score over 0.8

We then performed functional enrichment analysis using Gene Ontology (GO) terms, based on differentially expressed genes (DEGs) and differentially accessible promoter regions (DAPRs). We observed distinct regulatory landscapes across embryonic lineages (Figure 2D; Supplementary Tables 7 and 8). In the EPI, we identified a set of shared GO terms associated with neuronal development, morphogenesis, and cell differentiation, suggesting an early transcriptional priming toward ectodermal fates. Notably, several signaling pathways were marked by open promoter regions without corresponding gene expression, implying a poised yet transcriptionally inactive state. These include key regulators such as the ‘negative regulation of Notch signaling pathway’, ‘EGF receptor signaling pathway’, and ‘negative regulation of receptor signaling pathway via JAK-STAT’ (Supplementary Table 6), which may indicate epigenetic readiness for future activation. In the TE, GO enrichment predominantly highlighted terms related to energy production, molecular transport, and metabolic processes, consistent with the high biosynthetic and proliferative demands of this lineage. In contrast, in the HYPO, we observed a dynamic change in pathways and functions. During early hypoblast development (HYPO Ea), we observed enrichment for MAPK signaling-related terms, including ‘negative regulation of MAPK cascade’, ‘positive regulation of ERK1 and ERK2 cascade’, ‘regulation of JNK cascade’, and again the ‘negative regulation of Notch signaling pathway’. As development progressed to the late hypoblast stage (HYPO La), there was a noticeable shift toward enrichment of GO terms associated with BMP and TGF-β signaling, suggesting a transition in signaling pathway dominance between E7 and E11. This temporal shift may reflect functional maturation and lineage specification within the hypoblast.

### Comprehensive regulatory profiling of embryonic and extra-embryonic cell populations

We next explored the gene regulatory networks at play into those populations. To do this, we used SCENIC+^10^ to identify enhancer-driven regulons (eRegulons). These eRegulons represent functional gene modules regulated by enhancers. Each eRegulon is defined by a transcription factor (TF, which gives the eRegulon its name) and its associated set of target genes. Each target gene is co-expressed with the TF and has at least one accessible cis-regulatory element containing the TF’s DNA-binding motif. We identified 57 activator TFs whose target genes show both chromatin accessibility and induced expression, as well as 13 repressor TFs, characterized by closed chromatin at their target regions and suppressed gene expression (Supplementary Table 9). For each eRegulon, we visualised the expression levels of target genes (blue to red scale) and their chromatin accessibility (dot size) across clusters and developmental stages (Figure 4A). Compared to our previous SCENIC study on pig preimplantation embryos^5^, we identified fewer regulons but with higher confidence and 61% of TFs identified here were identified in both studies.

**Figure 4.**
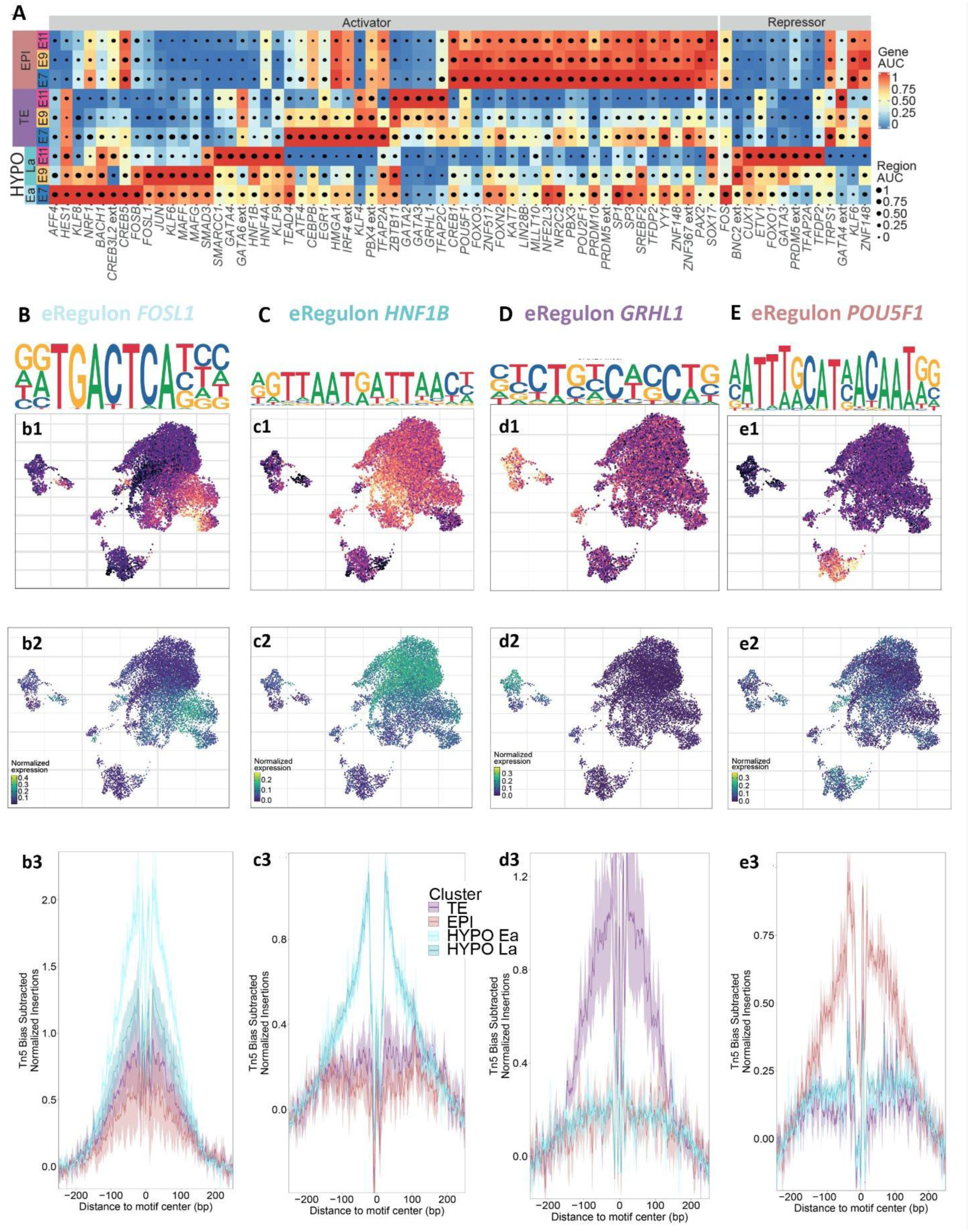
Active gene regulatory modules (eRegulon) of embryonic and extra-embryonic cell populations. **A.** Heatmap of eRegulons identified in each cell population and embryonic stage, classified either as activator or repressor. Dot size represents scaled region AUC and colour gradient represents target gene expression AUC. TFs name with ext (extended) represents TFs whose annotation is inferred based on motif similarity. **B-E**. Validation of eRegulons *FOSL1* (B), *HNF1B* (C), *GRHL1* (D) and *POU5F1* (E). **b1-e1.** UMAP visualisation of top motif enrichment in differentially accessible genes (DAGs) for *FOSL1, HNF1B, GRHL1* and *POU5F1* between each cluster. Gradient represents motif z-score calculated from ChromVar at the top motif PWMs used for the TFs. **b2-e2.** UMAP visualization of target gene expression levels for the eRegulons *FOSL1, HNF1B, GRHL1* and *POU5F1*. **b3-e3.** TFs footprint of *FOSL1, HNF1B, GRHL1* and *POU5F1* in each cluster at their known binding motifs. Footprints are inferred by assessing Tn5 activity bias around the DNA binding motif.

This analysis revealed key transcription factors with well-established roles in lineage specification, including *GATA2, GATA3*, and *TFAP2C* in TE, *GATA6, FOSL1*, and *HNF4A* in HYPO and *POU5F1, LIN28B*, and *ZNF148* in EPI. In addition, we identified new eRegulons, including *GRHL1, BNC2*, *IRF4*, *JUN*, *PAX2*, *PRDM5* and *TRPS1*. We also observed that some eRegulons regulated overlapping sets of genes due to their high motif similarity. These include the GATA TFs family (*GATA6, GATA4, GATA3*), TFs linked to the AP-1 complex (*ATF, JUN, BATF, FOS*)^13^ as well as two other families of TFs sharing a common set of genes: *MAFF*, *FOSL1*, and *BACH1* for the first set and *PRDM5*, *LIN28B*, and *PBX3* for the second set (Supplementary Figure 5).

To confirm the relevance of the identified eRegulons, we assessed motif enrichment within differentially accessible peaks (DAP) (Supplementary Table 10). For each cell cluster, we selected one of the most enriched motifs (*FOSL1* for Hypo Ea, *HNF1B* for Hypo La, *GRHL1* for TE, *POU5F1* for EPI) and visualised their enrichment scores on a UMAP projection (Figure 4b1-4e1). In parallel, we displayed the expression levels of their respective target genes (Figure 4b2–4e2). To further support the regulatory activity of these TFs, we inferred their footprint profiles around their respective DNA binding motifs by analysing Tn5 activity bias around their motif sequences (see Methods). The resulting TF footprints for FOSL1, HNF1B, GRHL1, and POU5F1 are shown in Figures 4b3–4e3. Together, these analyses confirm the high specificity of the identified eRegulons across distinct cell populations, reflected in both TF activity and target gene expression.

This analysis also revealed enriched motifs for TFs that have not been identified by our SCENIC+ analysis but are known to be lineage-specific markers, like *CDX*2 in TE or *MTA3* and *BCL11A* in EPI (Supplementary Figure 6).

Our analysis shows that the activity of eRegulons detected in the epiblast from E7 to E11 remained relatively stable, with some nuances, since most of them showed a slight decrease in activity from E7 to E11 (such as *POU5F1*), whereas *SOX17* and *TFAP2A* showed a slight increase in activity between E7 and E11. This sharply contrasts with extra-embryonic tissues (HYPO and TE) where eRegulon activity drastically changed between E7 and E11. Our analysis shows that this regulatory dynamic occurs concomitantly with the drastic morphological changes that occur in these tissues at the start of elongation, and should have an impact on the biology of these cells in terms of differentiation and cellular functions.

To understand whether this regulatory dynamic is associated with distinct biological functions, we performed GO and KEGG enrichment analyses on the target genes of each eRegulon, to identify relevant biological processes and pathways (Supplementary Tables 11-13). Surprisingly, most eRegulons are enriched for genes with functions specific to the biology of each cell population. In the EPI cluster, for instance, eRegulons predominantly target genes involved in embryonic development, particularly neural tissue formation (e.g., *LIN28B, PBX3, PRDM5, FOXO3*), as well as in the regulation of nuclear genome activity and DNA repair (*POU2F1, SP3, YY1, ZNF148, ZNF367, FOXN2*). Several eRegulons also control genes related to key signaling pathways, including Wnt (*PRDM5, LIN28B*), Sonic Hedgehog (*LIN28B*), RAP1 (*PAX2*), and Activin (*CREB1*). Additionally, some eRegulons appear more specialized, such as *MLLT10* regarding cell cycle regulation, *SREBF2* for cell metabolism, and *SP1* for intracellular trafficking (Figure 5, Supplementary Table 11).

In the TE, at early stages (E7 and E9), eRegulons are predominantly associated with terms linked to fundamental biological processes, like developmental processes (*KLF4, CEBP*), gene regulation (*KLF4*) and energy production (HMGA1). By E11, however, active eRegulons are associated with terms that shift towards functions related to more specialized cell functions, including lysosomes biology (*GATA3, GRHL1, TFAP2C*), transporter activity (*TFAP2C*), metabolism (*GRHL1, ZBTB11, GATA3*) and protein translation (*ZBTB11*) (Figure 5, Supplementary Figure 12).

A similar dynamic is observed during HYPO development, where eRegulon activity reflects cellular transitions. In HYPO Ea, active eRegulons are mainly associated with terms linked to morphogenesis and associated signaling pathways (*JUN, KLF6, MAFF*) that mirror HYPO formation. In contrast, eRegulons active in HYPO La are predominantly linked to cell metabolism and protein glycosylation (*SMARCC1, GATA4, HNF4A*) (Figure 5, Supplementary Table 14).

**Table 1.**
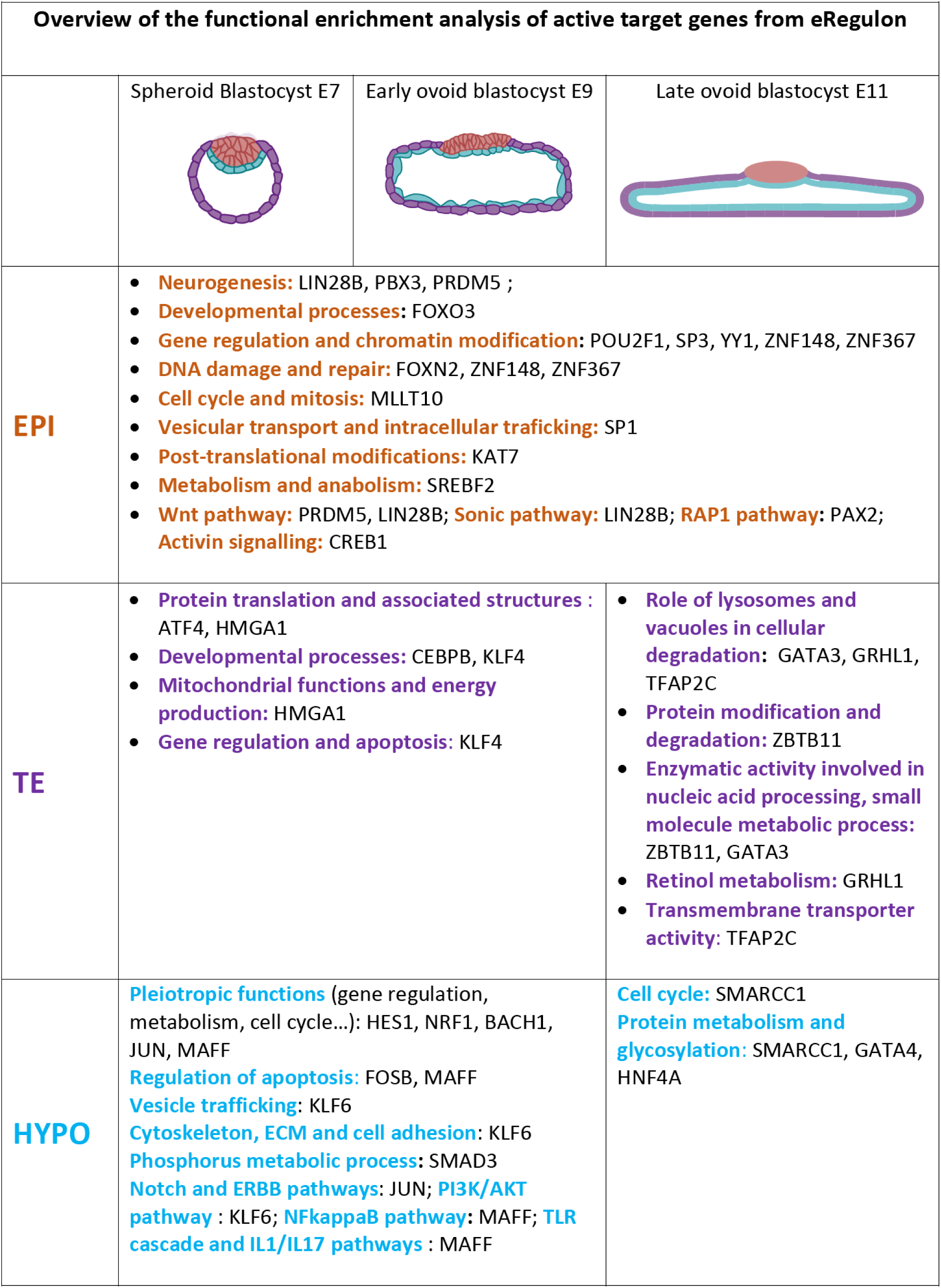
Summary of the functional enrichment analysis of active target genes inferred from eRegulon analysis across embryonic lineages in porcine blastocysts at days E7, E9 and E11. Each functional category is associated with eRegulons Transcription Factors. Colored text highlights distinct functional themes and signaling pathways identified within each lineage.

To complete our analysis on the regulatory landscape and its association with biological functions, we investigated peak-to-gene associations to identify potential cis-regulatory elements driving gene expression in each cell population. A total of 15,991 peak-to-gene regulatory links were identified (see Methods and Supplementary Table 6). To visualize these regulatory interactions, we selected links exhibiting a variance score above 0.25 and grouped the corresponding peaks into five different k-means clusters, representing the four main cell populations (Figure 6). For each cluster, the top 1,000 highly regulated genes (HRGs) with the highest number of associated peaks were subjected to functional enrichment analysis (Figure 6, Supplementary Table 14).

**Figure 6.**
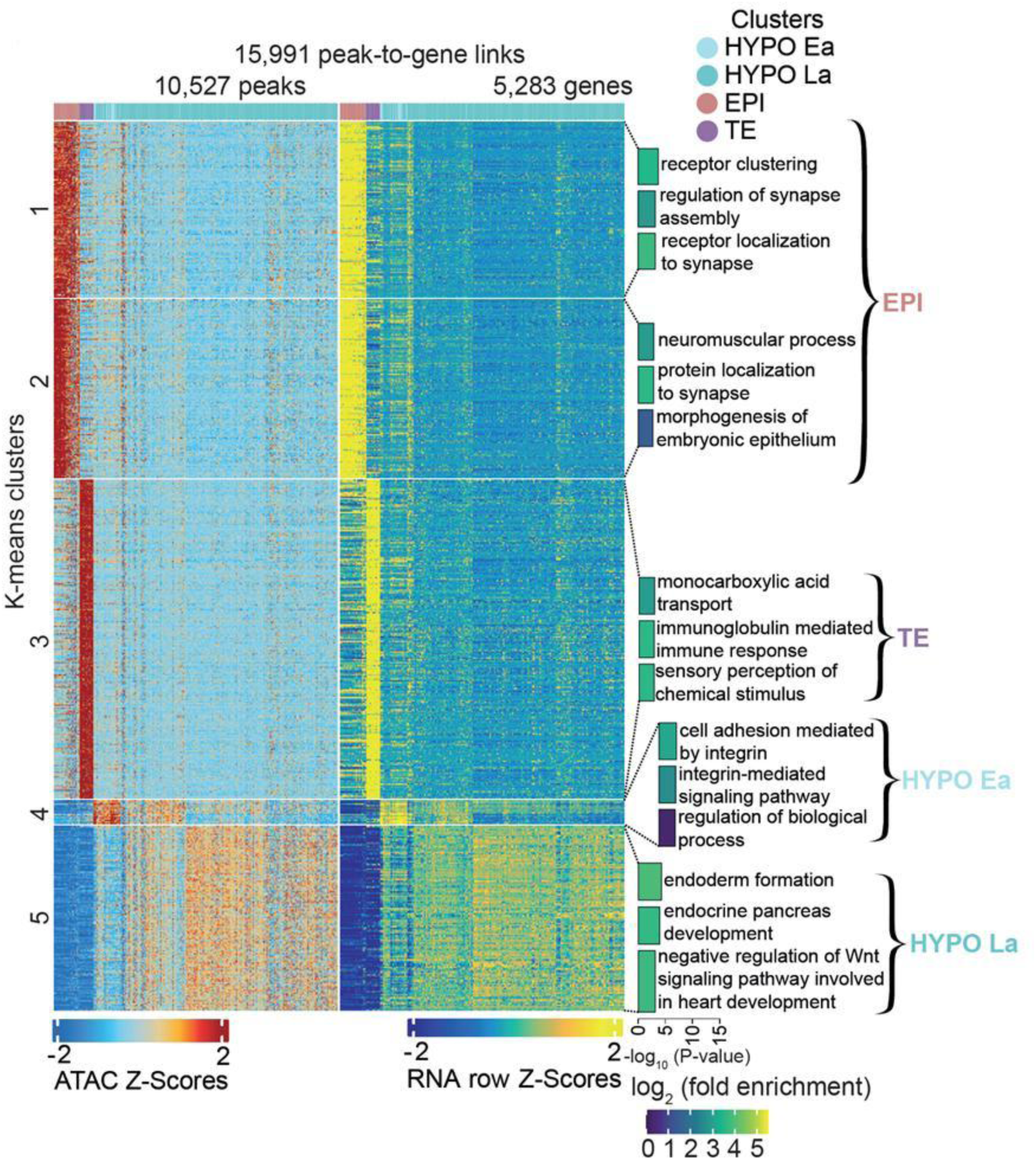
Heatmap representing chromatin accessibility on the left and gene expression on the right for 5,000 peak-to-gene linkages and clustered using the k-means clustering algorithms. In each cluster, the 1,000 most highly regulated genes (e.g. which have a high number of peaks linked to them) were used for Gene Ontology: Biological Process enrichment and the top 3 terms for each cluster are shown. Colour gradient represents the log of the fold enrichment among genes in peak-to-gene linkage.

This analysis reinforces the Gene Ontology (GO) annotations previously performed on differentially expressed genes (DEGs) and genes from active eRegulons, as shown in Figure 1G and Figure 5. The first two clusters of HRGs, predominantly expressed in EPI, are enriched for GO terms related to neurogenesis and morphogenesis. The third HRG cluster, specific to the TE, is enriched for pathways involved in lipid metabolism and transport. Additionally, this cluster also shows enrichment for GO terms related to immune response processes (Figure 6). In the fourth cluster, specific to hypoblast (HYPO), HRGs are associated with terms linked to cell adhesion processes mediated by integrins, while HRGs from cluster 5 genes are associated with terms linked to endoderm development and negative regulation of Wnt signaling pathways. Notably, among these HRGs, GATA6—a key transcriptional regulator of hypoblast lineage specification—was found to be associated with two distal enhancers strongly correlated to its expression in HYPO (asterisks in Supplementary Figure 7), mirroring similar enhancer–gene associations observed for GATA3 in TE and SOX2 in EPI (Figure 3).

## Discussion

This study provides the first comprehensive annotation of the regulatory landscape in the pig blastocyst, characterizing each distinct cell population by linking gene expression with chromatin accessibility.

This study completes our knowledge of the dynamics of the epigenome and gene regulation in mice and humans^14,15^ by focusing on a developmental stage that is specific to many other mammals, where blastocyst development continues for several days prior to implantation^1^. Our analyses enable us to annotate numerous regulatory elements whose accessibility correlates with the expression of known transcription factors for the different cell populations that compose the blastocyst (HYPO, EPI, TE). These results will support comparative studies on gene regulatory networks — both conserved and divergent — across mammalian species, with potential insights into species-specific traits and associated evolutionary strategies.

Our findings reveal two distinct regulatory dynamics between embryonic and extraembryonic tissues. In the epiblast, despite the divergence of preimplantation development in pigs from that of rodents and primates beyond the hatched blastocyst stage, the core regulatory gene networks, including *POU5F1, LIN28B, PRDM5, YY1, POU2F1*^11,16–19^, are conserved and show limited variation between embryonic day 7 and day 11. This also includes regulators of known signaling pathways involved in pluripotency, such as CREB1, which regulates the MEK/ERK pathway^20^, FOXO3, which regulates the PI3K/Akt^21^ and ZNF148, a Notch1 repressor^22^. These observations suggest that epiblast pluripotency is relatively stable during this 4-day window and resembles the pluripotent state described in other mammals. Given that gastrulation in pigs begins around embryonic day 11.5 at the tubular stage^6^, it is likely that major changes in gene regulatory networks and epigenomic states occur only at this tubular blastocyst stage, just beyond the ovoid stage examined in this study.

In the epiblast, among the identified eRegulons, several are linked to neural tissue development and neurogenesis, functions also found in the functional annotations of the highly regulated genes (HRGs). This supports previous observations in mice, where mouse embryonic stem cells (mESCs) were shown to exhibit a default proneural state in the absence of extrinsic signals^23^. Interestingly, we identified *MLLT10* as an eRegulon involved in cell cycle regulation, confirming prior findings in which *DOT1*, a known partner of *MLLT10*, plays a role in controlling the cell cycle and in the differentiation of both mouse and human pluripotent stem cells^24–26^. We also identified *SREBF2* as an eRegulon, with its target genes primarily associated with the regulation of cellular metabolism, particularly lipid metabolism. Its role in pluripotent stem cell biology remains largely unexplored.

In contrast, the regulatory dynamics in extraembryonic tissues are markedly different. We observed a rapid evolution in eRegulon activity during the studied time window, coinciding with the onset of extraembryonic tissue elongation. In both the trophectoderm (TE) and hypoblast (HYPO), certain eRegulons are predominantly active in the hatched blastocyst prior to elongation, while others become specifically activated during the initiation of the elongation process.

Regarding the hypoblast, we identified in this dataset two cell clusters specific to the early (HYPO Ea) and late (HYPO La) hypoblast, that we previously described^5^. In HYPO Ea, numerous eRegulons from the AP-1 transcription factor complex, including *FOSL1*, *BACH1*, *FOSB*, and *JUN*^27^, are associated with cell proliferation. Other eRegulons active in HYPO Ea, such as *KLF8*, *NRF1*, *CREB*, *KLF6*, and *MAFF*, have been previously identified in the visceral hypoblast subpopulation^5^. While most of these eRegulons function as pleiotropic regulators, some exhibit more specialized roles. For instance, *KLF6* predominantly regulates target genes involved in cell adhesion, extracellular matrix interactions, and vesicle formation and trafficking, whereas *FOSB* primarily controls the expression of genes linked to apoptosis.

In contrast, eRegulons associated with classical hypoblast markers from the *GATA* and *HNF* families, as well as *SOX17*^28^ show increased activity in the late hypoblast. This transition is accompanied by extensive remodeling of chromatin accessibility and enhancer activity, leading to a significant upregulation of the target genes of these eRegulons despite the fact that these transcription factors, such as *GATA4* and *GATA6*, are already expressed earlier as markers of hypoblast identity. In line with hypoblast maturation, we observed biological functions such as protein metabolism and glycosylation, which are characteristic of more differentiated hypoblast cells. These functions are absent in the early hypoblast, even though the transcription factors that drive them — including *SMARCC1*, *GATA4*, and *HNF4A* — are already expressed in HYPO Ea. This temporal shift between TFs expression and target gene activation, driven by differential chromatin accessibility, has also been observed in the mouse embryo, notably for *GATA3* in the trophectoderm^14^.

For the trophectoderm, although we were not able to identify subpopulations as in our previous study, we observed a strong dynamic in eRegulon activity across developmental stages, similar to what we found for the hypoblast. In the earlier, less mature stages, we identified eRegulons associated with morphogenesis (such as KLF4 and CEBPB) and mitochondrial activity (such as HMGA1). These findings support the role of KLF4 in trophoblast lineage specification, consistent with its previously established function as a key factor in reprogramming human fibroblasts into trophoblast stem cells^29^. Likewise, HMGA1 has been shown to regulate cell proliferation and migration in human trophectoderm^30^, and is itself regulated by the LIN28/let-7 axis, which plays a role in trophectoderm elongation in sheep^31^.

At later stages, we observed the activation of eRegulons corresponding to well-established TE-specific transcription factors, including *GATA2, GATA3*, and *TFAP2A/C*, as well as novel transcription factors and regulons not previously described in porcine TE, such as *GRHL1* and *ZBTB11*. Similar to what was observed for the hypoblast, there is a temporal shift between the activation of these eRegulons and the expression of their associated transcription factors^14^. This shift reflects rapid epigenomic remodeling and changes in the accessibility of regulatory regions in the target genes comprising these eRegulons across the different developmental stages. The functions associated with these eRegulons are characteristic of more differentiated cells and include lysosomal trafficking (GATA3, GRHL1, TFAP2C), post-translational modifications (ZBTB11), and membrane transporter activity (TFAP2C). GRHL1 has been shown to regulate trophoblast differentiation^32^ and has been proposed to act as a pioneer transcription factor that primes epithelial enhancers^33^. Interestingly, GRHL1 is itself regulated by KLF4, another transcription factor active in TE at earlier stages^34^.

Finally, as part of the Functional Annotation of ANimal Genomes (FAANG) international initiative, this work contributes to the FAANGSingleCell objectives by delivering single-cell atlases for key tissues in farm animal species^35^. By integrating gene expression and chromatin accessibility at single-cell resolution, the study robustly identifies key regulatory networks active during early developmental stages and will help to pinpoint genetic variants that may influence these regulatory mechanisms, with potential implications for traits of agricultural interest. To ensure the reproducibility and reusability of our datasets, all metadata and data generated in this study have been deposited in the FAANG Data Portal (https://data.faang.org), facilitating access and use by the broader research community.

## Conclusions

Altogether, our work using multiomic data at the single-cell level enabled us to identify changes in gene expression associated with epigenetic modifications during the preimplantation period of pig embryos. These changes are indicative of the maturation of the extra-embryonic tissues during elongation and are accompanied by a dynamic evolution of GRNs with decreasing or increasing activity of TFs in the different populations.

## Acknowledgments

We are grateful to the Genotoul bioinformatics platform Toulouse Occitanie (Bioinfo Genotoul, https://doi.org/10.15454/1.5572369328961167E12) for providing help, computing, and storage resources. We are grateful to people from the INRAE experimental farm GenESI (https://doi.org/10.15454/1.5572415481185847E12) who took care of the animals. We are grateful to Sylvain Bourgeois and the people from the Unité Expérimentale de Physiologie Animale de l’Orfrasière (UEPAO) INRAE. We thank the @BRIDGe facility for excellent technical assistance. We thank people from our respective laboratories for their helpful comments and discussion. We thank the foundation Vaincre le Cancer-NRB for the acquisition of the Chromium 10X machine.

## Methods

### Embryo Production

All embryos were produced at the INRAE experimental unit GenESI (Rouillé, France). Metadata related to the biological samples used in this study have been submitted to FAANG Data Portal (https://data.faang.org/home) and BioSamples (https://www.ebi.ac.uk/biosamples/), and are summarised in Supplementary Table 1.

The oestrous cycle of each sow was synchronised using Altrenogest (Regumate), a synthetic progestin, for 18 days. The day after, sows were artificially inseminated, and the insemination was repeated the following day. When the gestational time matched the embryonic stage to be sampled (7, 9, and 11 days post-insemination), the sows were transported from the breeding unit (Rouillé, France) to the slaughterhouse (Nouzilly, France). They were stunned by electronarcosis and bled. The uterus was clamped and rapidly extracted from the abdominal cavity. Then, embryos were collected into two 50 ml tubes by retro-flushing of the uterine horns from the bottom of the uterine horn upwards (ovary) in 100 ml of physiological saline solution. The detailed protocol is publicly accessible on the FAANG Data Portal: https://api.faang.org/files/protocols/samples/INRAE_SOP_PLUS4PIGS_EMBRYOS_SAMPLING_PROTO2_20230131.pdf.

### Preparation of Single Nuclei Suspension

Once recovered, embryos were staged, pooled, and transported in DMEM/F12 medium (E9 and E11) or IMV Embryo holding media (E7) to the molecular biology laboratory in a thermostatically controlled chamber at 38°C. Upon arrival, embryos were transferred to a 4-well dish and staged again under a stereomicroscope. When necessary, embryos of the same stage were pooled together into a drop of DMEM/F12 or IMV Embryo holding media and processed for cell dissociation. The full protocol is publicly accessible on the FAANG Data Portal: https://api.faang.org/files/protocols/samples/INRAE_SOP_PLUS4PIGS_EMBRYOS_DISSOCIATION_PROTO3_20230131.pdf. All embryos (E7 to E11) were incubated in pre-warmed Accutase for 10 minutes, then in pre-warmed TrypLE for 10 minutes, followed by mechanical dissociation by repeated pipette aspiration.

Nuclei were isolated following the 10X Genomics protocol: CG000365 • Rev A Nuclei Isolation for Single Cell Multiome. Nuclei for embryonic cells were isolated following the same protocol from 10X Genomics, using the dedicated protocol for low cell input nuclei isolation (Appendix section). For E9 and E11 embryos, around 40,000 cells were used for each preparation. For E7 embryos, all cells (∼7,000) were used for one reaction. Each single cell suspension was washed twice with 50μl of ice-cold PBS containing 0.04% BSA and placed in a 0.2ml tube. Cells were centrifuged for 1 minute at full speed using a labtop minifuge. After the second wash, the supernatant was removed and 45μl of ice-cold lysis buffer was added (10mM Tris-HCl pH7.4; 10mM NaCl; 3mM MgCl2; 0.1% Tween-20; 0.1% Nonidet P40; 0.5% BSA; 1mM DTT; 0.5U/μl RNase inhibitor; 0.01% digitonin). Cells were resuspended by pipetting up and down 3 times and were incubated on ice for exactly 3’30. Then, 50μl of ice-cold Wash Buffer (10mM Tris-HCl pH7.4; 10mM NaCl; 3mM MgCl2; 0.1% Tween-20; 0.5% BSA; 1mM DTT; 0.5U/μl RNase inhibitor) was added carefully to the tube without disrupting the pellet. Cells were again centrifuged 1 minute at full speed using a labtop minifuge. 95μl of supernatant was then removed without disrupting the nuclei pellet and 95μl of chilled Diluted Nuclei Buffer was carefully added to the tube. Cells were again centrifuged 1 minute at full speed using the labtop minifuge. The supernatant was removed without touching the bottom of the tube to avoid dislodging the nuclei pellet. The nuclei pellet was then resuspended in 7μl of chilled Diluted Nuclei Buffer.

To quantify nuclei, 2μl of nuclei suspension were mixed with 8μl of Diluted Nuclei Buffer and 10μl of Trypan Blue and counted using a Malassez cell counting chamber. Around 5,000 nuclei/μl have been recovered from each prep. 25,000 nuclei resuspended in 5μl of Nuclei Dilution Buffer are processed for transposition according to: ChromiumNextGEM_Multiome_ATAC_GEX_User_Guide_RevB.

The full protocol is publicly accessible on the FAANG Data Portal: https://api.faang.org/files/protocols/samples/INRAE_SOP_PLUS4PIGS_.pdf.

### scMultiome library preparation and sequencing

Libraries were sequenced on an Illumina NovaSeq6000 and subsequently aligned to the *Sus scrofa* genome (version 11.1). Gene positions were annotated according to Ensembl build 108 and filtered based on biotype (only protein-coding, long intergenic non-coding RNA, antisense, immunoglobulin or T-cell receptor). Following this, CellRanger ARC (version 2.0.0) was employed to generate the count matrices based on this annotation.

### ArchR gene and genome annotation

The data were processed using the ArchR R package (1.0.3)^36^. Gene annotation parameters were developed using the GTF files produced earlier and the Org.Ss.eg.db package. Genome annotation was conducted similarly, applying filters to exclude contigs that were not detected in either the ATACseq or the RNAseq datasets (https://doi.org/10.5281/zenodo.10844556), while also including the mitochondrial chromosome. Due to the presence of short contigs, the downstream and upstream extensions for the gene score matrix function were set to a minimum of 0 and a maximum of 100,000.

### Single-cell multi-omics analysis on the embryos

A quality control was applied and cells were excluded if they had more than 1e+10 or fewer than 1,000 fragments and a minimum enrichment of Transcription Start Site (TSS) of 4. Cells were retained only if they were present in both ATACseq and RNAseq assays. Subsequently, we filtered out cell doublets using ArchR functions addDoubletScores and filterDoublets with default parameters, resulting in a total of 12,778 cells (Supplementary Table 2). After filtering, we generated Latent Semantic Index (LSI) for both assays.

For the ATACseq part, we used the TileMatrix with the following parameters: a resolution of 2, a sample size of 30,000 cells, n.start set to 10 and the number of fragments as coverage. For the RNAseq part, we used the GeneExpressionMatrix with the following parameters: a resolution of 2, a sample size of 30,000 cells, n.start set to 10 and the number of fragments as coverage. We used the number of Unique Molecular Identifiers (UMI) as coverage, selected the 2,500 most variable genes and deactivated the binarization. A new dimensional reduction was then created using the two previously generated LSI with the addCombinedDims function.

Batch corrections for each sample were applied using the Harmony implementation in ArchR on the combined dimensional reduction. A UMAP was generated on the corrected dimensions reduction using 15 neighbors, a minimal distance of 0.8 and a cosine metric. Then a first clustering was conducted at a resolution of 0.1 (Figure 1B & C).

Group coverage and peak identification were conducted using ArchR function addGroupCoverage and addReproductiblePeakSet, alongside MACS (v2.2.7.1)^37^ with a genome size of 1,341,049,888.

Peak enrichment analyses were performed using the getMarkerFeatures function, utilizing the wilcoxon method along with correction for TSSEnrichment and log10 bias. Additionally, connections between peak and gene expression were established using the addPeak2GeneLinks function in ArchR, applying Harmony^38^ for dimensionality reduction and the gene expression matrix. The resulting data was incorporated into the differential peak table (supplementary Table 6) when significant correlations were identified.

### Cluster assignment

To perform embryo assignment (Supplementary Figure 2), we used our previous scRNAseq data from pig embryos^5^. Cell mapping was conducted using ArchR addGeneIntegrationMatrix function with the following parameters: dimensional reduction was carried out using Harmony, cell sample size of 50,000, number of dimensions set to 60 (utilizing all available dimensions), 4,000 variables genes, the cca reduction. Subsequently, we applied the FindAnchorFunction from ArchR with the following parameters: k.anchor = 80, k.filter = 1600, k.score = 200, max.features = 800, and n.trees = 800. Based on these results, we confidently identified clusters of epiblast and trophectoderm. Additionally, we distinguished two clusters of hypoblasts (early and late), though some cells within the late hypoblast cluster displayed a lower confidence score. All identified clusters expressed specific gene markers indicative of their respective populations^3,4^.

### Peaks assignation for genomic features

Promoter’s maps were generated using ArchR classification of peaks. Peaks identified with ArchR were classified into four categories: introns, exons, distal and promoter for the peak situated within 2,000 bp upstream and 100 bp downstream of the TSS site.

### SCENIC+ analysis

To run SCENIC+^10^ on the pig genome, we created a custom database. Transcription factor lists were generated following the methodology used for AnimalTFDB 4.0^39^, utilizing the pig Ensembl build 108. The motif-to-transcription factor annotation was adapted to pig by leveraging genes orthology through gprofiler2^40^. The motif database was constructed using scripts from aertslab (github.com/aertslab/create_cisTarget_databases) focusing on the best transcript score for each region of the genomes based on the 10 kb upstream regions.

The pipeline was executed on embryo using a combination of clusters and stages as cell aggregations, excluding categories with fewer than 20 cells (specifically, Cluster Hypo Ea at E7 and E9, and Cluster HYPO La at E7), applying default parameters and adapting annotations for the pig genome. For CisTopic analysis, we retained 16 topics. Following the SCENIC+ results, chromatin and transcriptome scores were calculated using the score_eRegulons function with an auc_threshold set to 0.05. We filtered the resulting regulons by rejecting extended regulons if direct regulons already existed, excluding regulons with fewer than 10 genes and those where region accessibility correlated with transcriptomics.

### Motif & Footprints analysis

Motif analysis was conducted using the aertslab v10 motif collection^10^. Motif annotations were incorporated into the ArchrProject through the addMotifAnnotations function, with parameters set to a cutoff of 5e-05, a width of 7 and utilizing version 2. Motif enrichment was subsequently performed using the peakAnnoEnrichment function from ArchR, with a filter applied for a minimal false discovery rate (FDR) of 0.1 and a log-fold greater than 0.5. The background peak and Deviations Matrix were computed using the addDeviationsMatrix and addBgdPeaks functions from ArchR, employing default parameters.

After obtaining the motif enrichment results, the motif positions were acquired using ArchR getPositions function with default settings. Footprints were generated using the ArchR getFootprints function, and these footprints were visualized through the plotFootpints function, utilizing “Substract” normalization method and a smoothing window that corresponds to the motif length.

### Functional enrichment

Functional enrichment of gene expression between clusters (Figure 2D) was assessed using marker genes from RNAseq, which were obtained via the ArchR getMarkerFeatures function on the GeneExpressionMatrix. The Wilcoxon test was employed, along with a correction for the number of UMIs. Following this, an enrichment analysis was conducted using the gprofiler2 package for all identified genes, focusing on the *Sus scrofa* organism and applying default parameters to the 2,000 most enriched genes.

Similarly, functional enrichment of peak differential accessibility between clusters was determined by linking genes to the corresponding peaks or if no gene was associated, using the nearest gene in the genome. This was also achieved using ArchR getMarkerFeatures on GeneExpressionMatrix using the Wilcoxon test with a correction for the number of UMIs. The resulting gene table underwent an enrichment analysis with the gprofiler2 R package for the *Sus scrofa* organism, using default parameters on the 2,000 most enriched genes.

For peak-to-gene heatmap functional annotations (Figure 6), we implemented methods outlined in GreenLeaf et al^41^. We selected the 1,000 most frequent genes for each k-means cluster, then gene ontology (GO) enrichment was calculated using the TopGO package^42^. The runTest function was applied using GO with specific parameters: Biological Process, mapping to the *Sus scrofa* database via symbol gene names, a minimal threshold of 5 genes, the weight01 algorithm and the Fisher test. Subsequently, the GenTable function was utilized to select the 50 top GO terms.

## Data availability

All raw sequencing data are available in FAANG Data portal (https://data.faang.org/home) under accessions PRJEB81663 and PRJEB83269.

## Code availability

The code used for the analysis is available at https://github.com/INSERM-U1141-Neurodiderot/pig_embryo_multiome. Txdb package used for Archr is available at https://doi.org/10.5281/zenodo.10844555. Pig SCENIC+ database is available at https://doi.org/10.5281/zenodo.14833685.

## Supplementary Figures

**Supplementary Figure 1.**
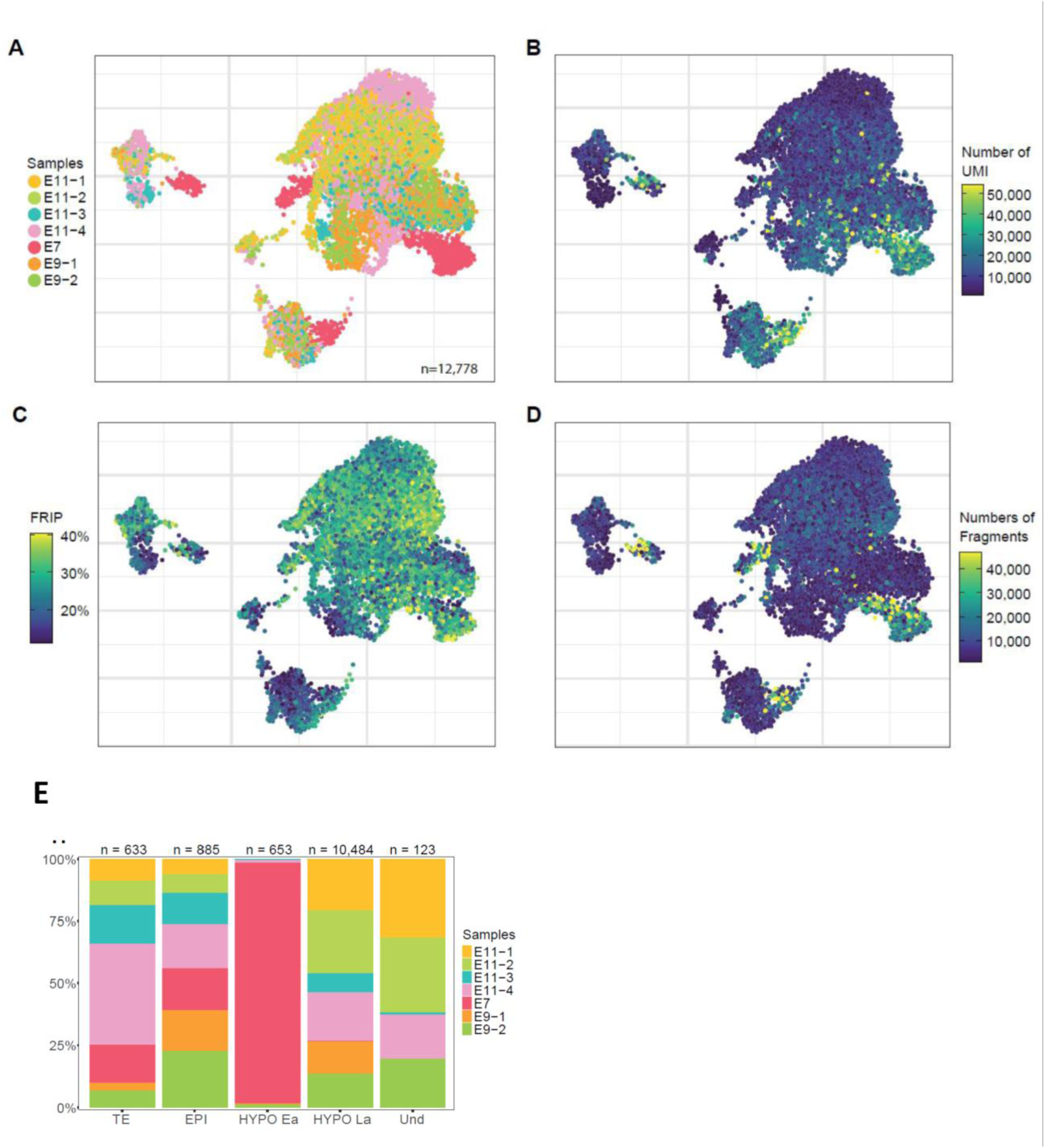
Visualization of quality control metrics from embryo scMultiome sequencing. **A.** UMAP representation of all single-cell passing quality control, coloured by the sample of origin (embryos from E7 to E11, see also Supplementary Table 2) **B**. UMAP representation of the number of RNA-seq UMI per cell. **C.** UMAP representation of the fraction of reads in peaks (FRIP) per cell. **D.** UMAP representation of the number of ATAC-seq reads fragments per cell. **E.** UMAP representation of identified cell clusters: EPI (red), TE (purple), HYPO Ea (light blue), HYPO La (dark blue) and Und (orange).

**Supplementary Figure 2.**
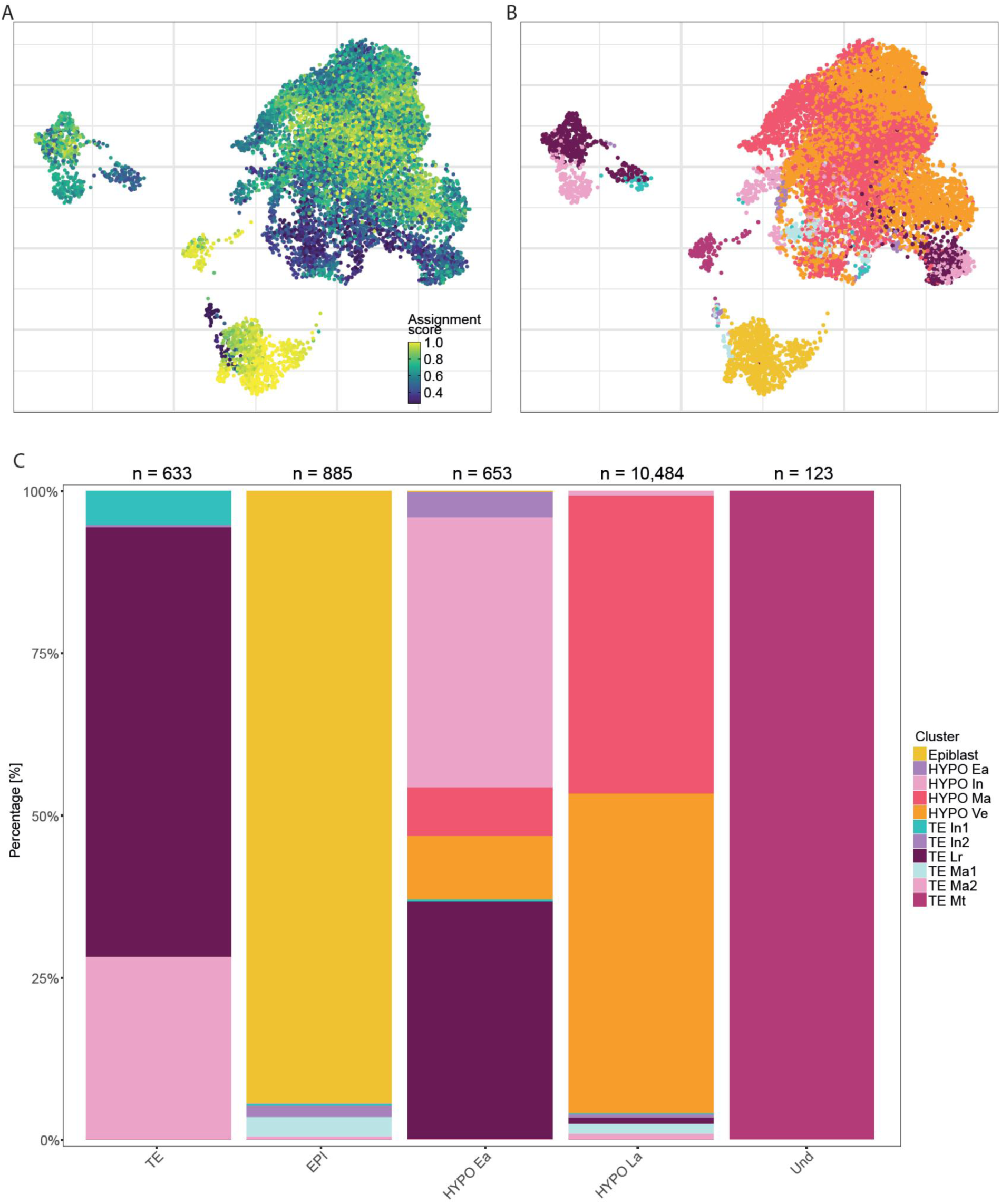
Annotation of Embryonic Cells Using a Pig Embryo scRNA-seq Reference Atlas. **A.** UMAP representation of the assignment score per cell. **B.** UMAP representation of the population assignment for each cell, with cluster colors corresponding to those in the main figure legend. **C.** Proportion of cells from each cluster assigned to the different populations.

**Supplementary Figure 3.**
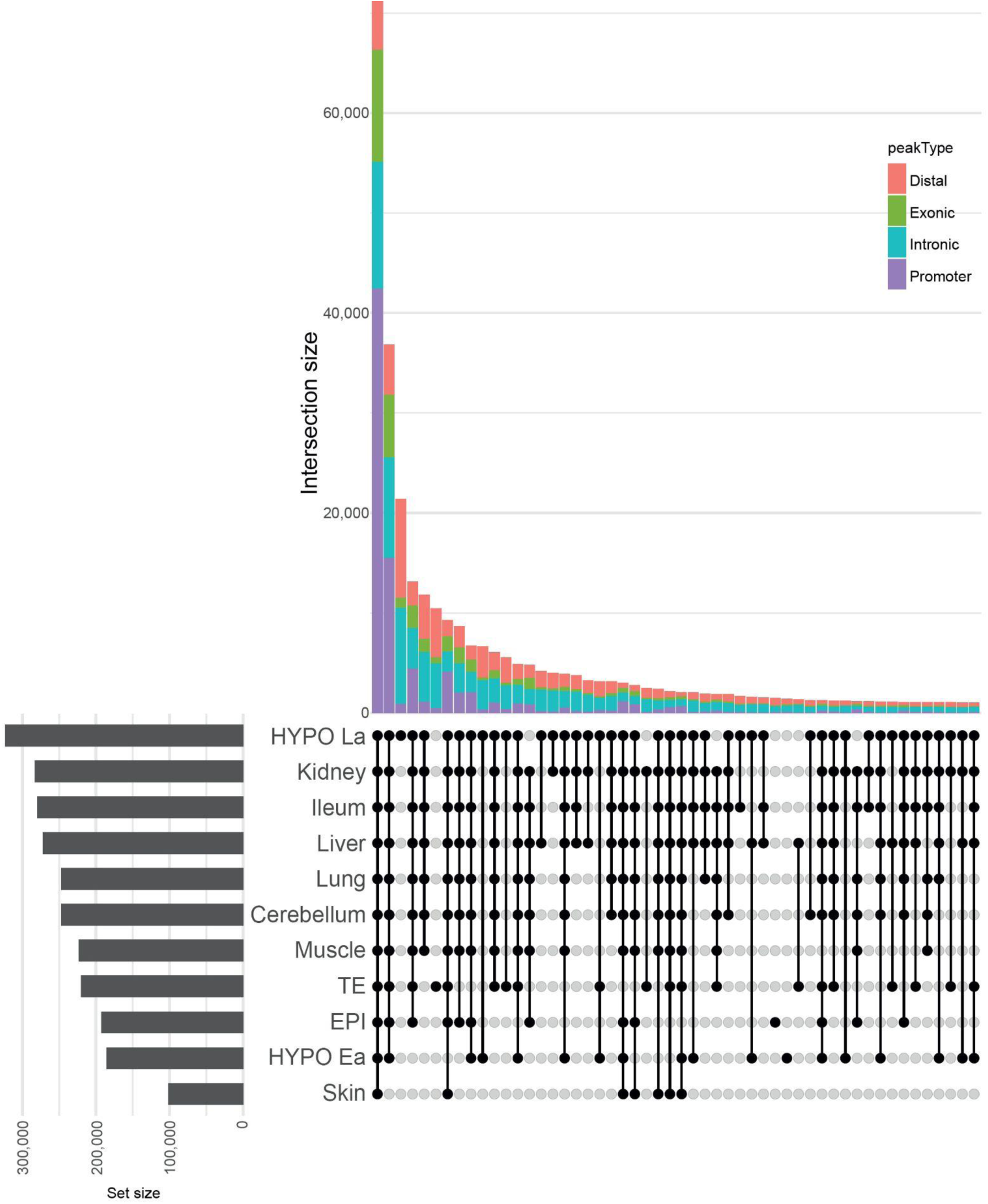
Upset plot comparing the number of peaks overlap between embryonic cell populations and somatic tissues.

**Supplementary Figure 4.**
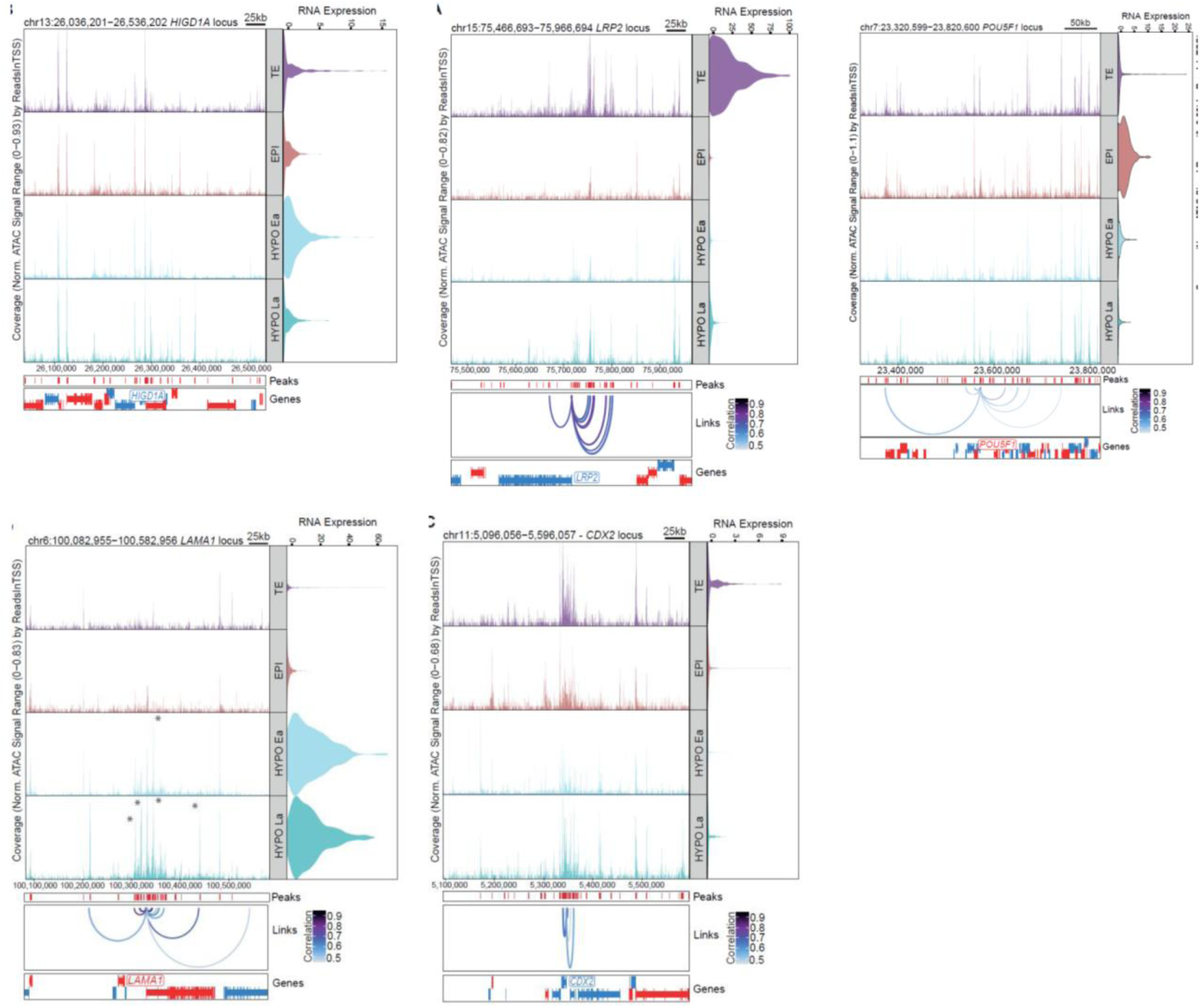
Peak-to-gene linkages for HYPO, TE and EPI markers. Genomic track for chromatin accessibility around the *HIGD1A, LAMA1, LRP2, CDX2* and *POU5F1* loci. Right: expression levels are shown in the violin plot for each cell type. Loops shown below the top panel indicate peak-to-gene linkages identified on the full dataset.

**Supplementary Figure 5.**
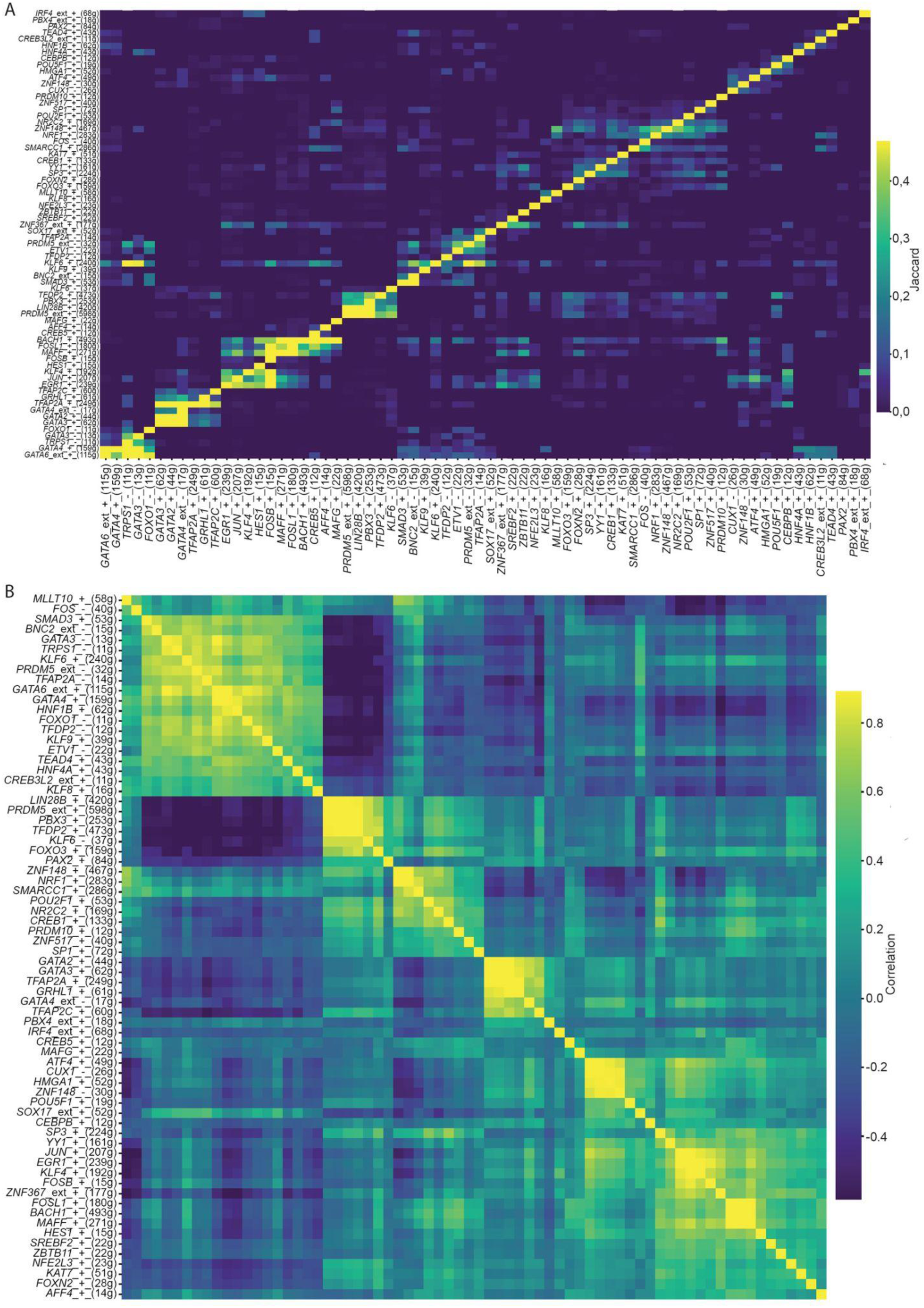
SCENIC+ similarity between eRegulons in the embryos. **A**. Heatmap showing the Jaccard similarity between eRegulons compositions **B.** Heatmap showing the correlation of motifs assignments associated with each eRegulons.

**Supplementary Figure 6.**
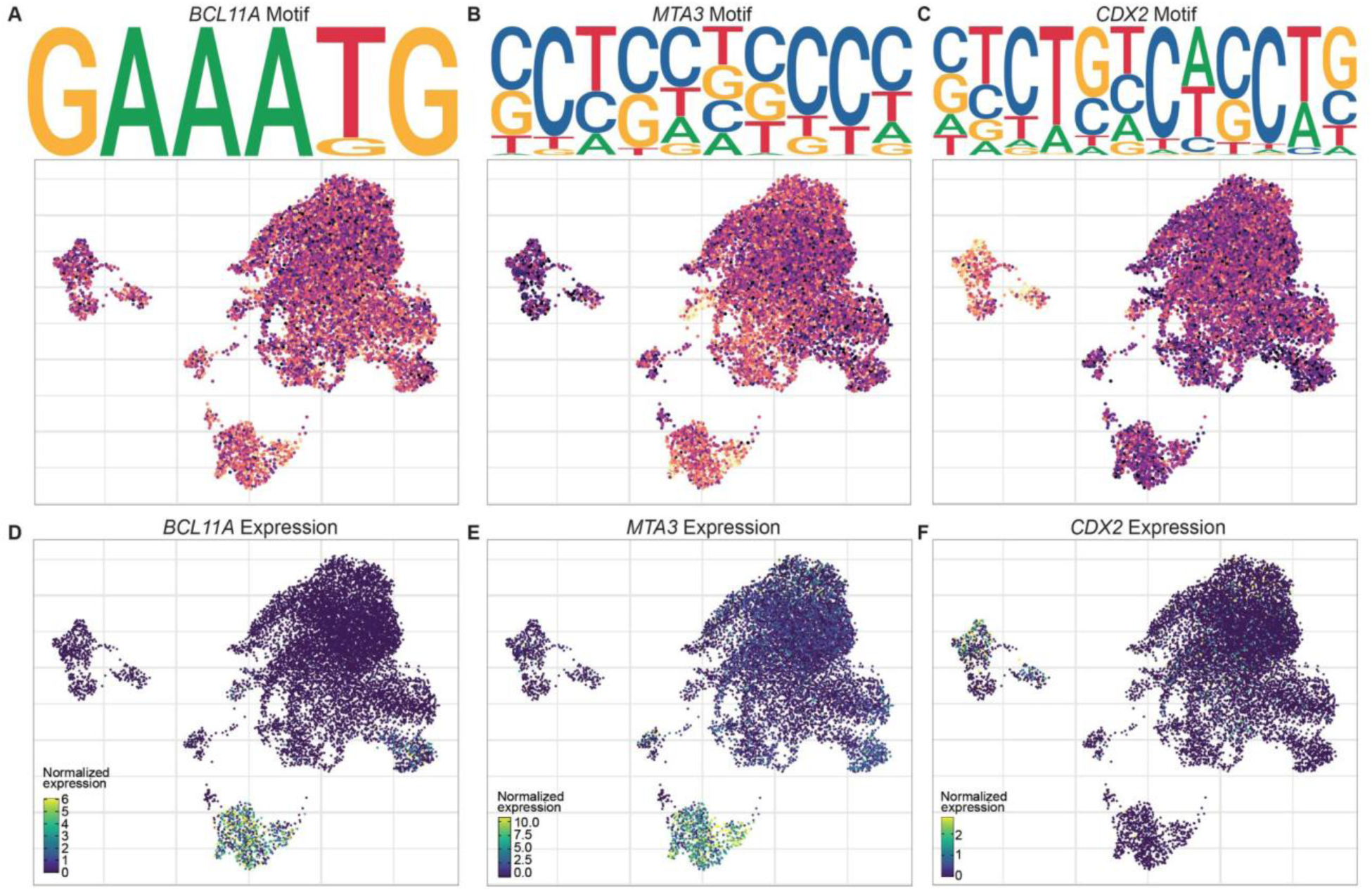
A. UMAP visualisation of *BCL11A* motif, colour gradient represents motif z-score calculated from ChromVar, at the top motif PWMs used for the TFs. **B.** Same as A for *MTA3* motif. **C.** Same as A for *CDX2* motif. **D.** UMAP visualisation of *BCL11A* expression, colour gradient represents normalised expression. **E.** Same as C for *MTA3* expression. **F.** Same as C for *CDX2* expression.

**Supplementary Figure 7.**
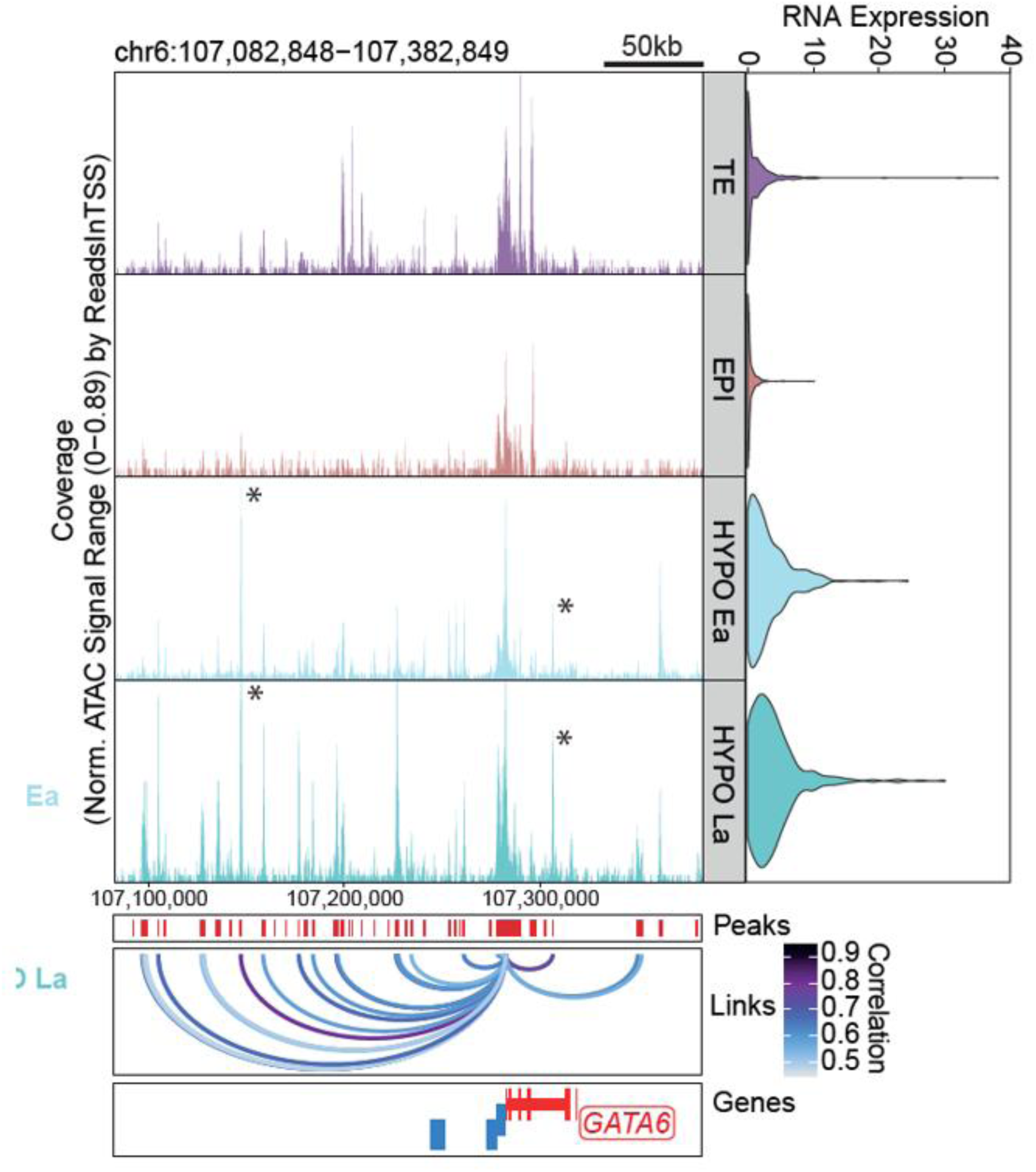
Profiles of peak-to-gene linkages for *GATA6* in HYPO, TE and EPI of pig embryos.

## Legends of supplementary tables

**Supplementary Table 1: Sample metadata.**

Table summarizing the characteristics of embryos used in this study, including: sow ID and breed, date of sampling, sow BioSample ID, boar ID and breed, embryo production protocol, developmental stage, number of embryos, BioSample names and IDs (for both embryos and single-cell specimens), stage descriptions, library names, and ENA accession IDs (studies and experiments).

**Supplementary Table 2: metrics of library sequencing and library qualities.**

Table summarizing sequencing depth and library quality metrics, including: number of raw RNA-seq reads, percentage of RNA-seq reads confidently mapped to the genome, median UMI counts per cell, number of detected genes, median high-quality fragments per cell, number of raw ATAC-seq reads, TSS enrichment score, number of confidently mapped ATAC-seq reads, number of cells before quality control, and number of cells retained after quality control.

**Supplementary Table 3: Intersection of DNA accessible peaks between somatic (Cerebellum, Ileum, Kidney, Liver, Lung, Muscle, Skin) and embryonic (TE, EPI, HYPO Ea, HYPO La) tissues.**

Table summarizing the intersection of peaks between tissues. These are the 11 tissues being compared. In the first column, each position in the binary string corresponds to one tissue in this exact order: `1` = peaks are overlapping in that tissue, `0` = peaks are not overlapping in that tissue.

**Supplementary Table 4: Differentially Accessible Peaks between cell clusters identified in this study using the getMarkers function from the ArchR package.**

Table summarizing differentially accessible peaks between cell clusters including: peakName, chromosome location, peak start, peak end, Log2FC, FDR, MeanDiff, Peak score, Nearest Gene, Peak location, P2G, P2G Correlation and P2G associated gene.

**Supplementary Table 5: Differentially Expressed Genes between cell clusters identified in this study using the getMarkers function from the ArchR package.**

Table summarizing differentially expressed genes between cell clusters including: gene name, Log2FC, FDR and MeanDiff.

**Supplementary Table 6: Peak-to-gene linkages in the embryos.**

Table summarizing the peak-to-gene linkages passing the correlation cutoff, including:gene-to-peak Name, chromosome location, start, end, Correlation value, FDR, VarQATAC, VarQRNA and Gene name.

**Supplementary Table 7: Functional enrichment analysis from upregulated genes in each cell cluster.**

Table of significantly enriched Gene Ontology (GO) terms across cell clusters. Columns include: GO term ID, GO term description, number of annotated genes, number of significant genes, expected, p-value.

**Supplementary Table 8: Functional enrichment analysis from genes associated with differentially accessible peaks.**

Table of significantly enriched Gene Ontology (GO) terms across cell clusters. Columns include: GO term ID, GO term description, number of annotated genes, number of significant genes, expected, p-value.

**Supplementary Table 9: Identified eRegulons in each cell cluster using SCENIC+.**

The first sheet provides detailed information for each eRegulon, including its target genes, associated peaks, and the relative importance of each feature for the corresponding transcription factor. In the two first columns, “+_+” means positive correlation of TF-to-gene and region-to-gene, corresponding to activators associated with opened chromatin at the target regions and induced gene expression of the target gene; *“*-_-” means negative correlation of TF-to-gene and region-to-gene, corresponding to repressors, associated with closed chromatin of the target regions and repressed expression of the target gene. The first two columns also indicate the number of target regions (first column) and of target genes (second column).

The second sheet provides information on the composition of each regulon, including associated peaks and target genes. In the first column, “+_+” means positive correlation of TF-to-gene and region-to-gene, corresponding to activators associated with opened chromatin at the target regions and induced gene expression of the target gene; *“*-_-” means negative correlation of TF-to-gene and region-to-gene, corresponding to repressors, associated with closed chromatin of the target regions and repressed expression of the target gene.

The third sheet indicates for each Regulon the reference of its associated binding motifs.

**Supplementary Table 10: Motif enrichment in peaks of accessible chromatin across cell clusters.**

Columns include: motif_group, motif ID, enrichment scores respectively in TE, EPI, HYPO Ea, HYPO La, maximum enrichment score, TF (transcription factors known to bind the motif). Motif collection and TF annotations come from Aertslab human motif database with orthology provided by g:Profiler.

**Supplementary Table 11: Functional enrichment analysis from genes of eRegulons active in EPI.**

Functional enrichment analysis of genes composing eRegulons was performed using gProfiler.

Each sheet represents one eRegulon, columns include: source (database), term_name, term_ID, adjusted p-value, term size, query_size, intersection_size, effective_domain_size and intersections (gene list).

**Supplementary Table 12: Functional enrichment analysis from genes of eRegulons active in TE.**

Functional enrichment analysis of genes composing eRegulons was performed using gProfiler.

Each sheet represents one eRegulon, columns include: source (database), term_name, term_ID, adjusted p-value, term size, query_size, intersection_size, effective_domain_size and intersections (gene list).

**Supplementary Table 13: Functional enrichment analysis from genes of eRegulons active in HYPO.**

Functional enrichment analysis of genes composing eRegulons was performed using gProfiler.

Each sheet represents one eRegulon, columns include: source (database), term_name, term_ID, adjusted p-value, term size, query_size, intersection_size, effective_domain_size and intersections (gene list).

**Supplementary Table 14: Enriched pathways from peak-to-gene linkages.**

Gene Ontology : Biological Process positively enriched in each clusters of peak-to-genes linkage

**Supplementary File 1: Annotation file of chromatin accessibility peaks from the whole dataset.**

The table includes: peakName, chromosome, start, end, score (measure of peak strength), replicateScoreQuantile (quantile rank of the score across replicates), groupScoreQuantile (quantile rank of the score across experimental groups), Reproducibility, distToGeneStart, nearestGene, peakType, distToTSS, nearestTSS, proportion of GC

