## Supplementary Table 1 for "Single-cell omics reveal distinct gene regulatory dynamics underpinning embryonic and extraembryonic lineage functions during pig blastocyst development"

| Sow (ID & breed) | Date of sampling | BioSample ID sow | Boar (ID & breed) | Protocol for embryo production | Stage | Number of embryos | BioSamples’ name (embryos & single cell specimen)  (prefix: INRAE_Plus4PigS_) | BioSamples’ IDs (embryos & single cell specimen) | Stage (description) | Library name :  (prefix : SSC_INRAE_Plus4PigS_ | Accession ID (ENA)  Studies :  PRJEB81663  PRJEB83269 |
| --- | --- | --- | --- | --- | --- | --- | --- | --- | --- | --- | --- |
| 4290  Large White | 19/05/2021 | SAMEA112465595 | 1245  Large White | normal breeding procedure | D7 | 12 | embryo_D7_8  embryo_D7_omic_pool_1 | SAMEA112465630  SAMEA115811553 | spheroid blastocyst | embryo_multiome_scRNAseq_D7_1  embryo_multiome_scATACseq_D7_1 | ERX13271546  ERX13271555  ERX13450981 |
| 4239  Large White | 19/05/2021 | SAMEA112465596 | 1245  Large White | normal breeding procedure | D7 | 11 | embryo_D7_10  embryo_D7_omic_pool_1 | SAMEA112465632  SAMEA115811553 | hatched blastocyst | embryo_multiome_scRNAseq_D7_1  embryo_multiome_scATACseq_D7_1 | ERX13271546  ERX13271555  ERX13450981 |
| 4220  Large White | 19/05/2021 | SAMEA112465597 | 1245  Large White | normal breeding procedure | D7 | 14 | embryo_D7_12  embryo_D7_omic_pool_1 | SAMEA112465634  SAMEA115811553 | hatched blastocyst | embryo_multiome_scRNAseq_D7_1  embryo_multiome_scATACseq_D7_1 | ERX13271546  ERX13271555  ERX13450981 |
| 4182  Large White | 19/05/2021 | SAMEA112465599 | 1245  Large White | normal breeding procedure | D7 | 9 | embryo_D7_14  embryo_D7_omic_pool_1 | SAMEA112465636  SAMEA115811553 | hatched blastocyst | embryo_multiome_scRNAseq_D7_1  embryo_multiome_scATACseq_D7_1 | ERX13271546  ERX13271555  ERX13450981 |
| 4289  Large White | 19/05/2021 | SAMEA112465600 | 1245  Large White | normal breeding procedure | D7 | 9 | embryo_D7_16  embryo_D7_omic_pool_1 | SAMEA112465638  SAMEA115811553 | hatched blastocyst | embryo_multiome_scRNAseq_D7_1  embryo_multiome_scATACseq_D7_1 | ERX13271546  ERX13271555  ERX13450981 |
| 4246  Large White | 19/05/2021 | SAMEA112465601 | 1245  Large White | normal breeding procedure | D7 | 8 | embryo_D7_18  embryo_D7_omic_pool_1 | SAMEA112465640  SAMEA115811553 | hatched blastocyst | embryo_multiome_scRNAseq_D7_1  embryo_multiome_scATACseq_D7_1 | ERX13271546  ERX13271555  ERX13450981 |
| 4181  Large White | 19/05/2021 | SAMEA112465602 | 1245  Large White | normal breeding procedure | D7 | 9 | embryo_D7_20  embryo_D7_omic_pool_1 | SAMEA112465642  SAMEA115811553 | hatched blastocyst | embryo_multiome_scRNAseq_D7_1  embryo_multiome_scATACseq_D7_1 | ERX13271546  ERX13271555  ERX13450981 |
| 4278  Large White | 03/06/2021 | SAMEA112465604 | 1000  Large White | normal breeding procedure | D9 | 3 | embryo_D9_7  embryo_D9_omic_pool_1 | SAMEA112465644  SAMEA115811554 | ovoid blastocyst 5mm | embryo_multiome_scRNAseq_D9_1  embryo_multiome_scATACseq_D9_1 | ERX13271547  ERX13271556  ERX13450982 |
| 4300  Large White | 03/06/2021 | SAMEA112465605 | 1000  Large White | normal breeding procedure | D9 | 22 | embryo_D9_10  embryo_D9_omic_pool_1 | SAMEA112465647  SAMEA115811554 | ovoid blastocyst 5mm | embryo_multiome_scRNAseq_D9_1  embryo_multiome_scATACseq_D9_1 | ERX13271547  ERX13271556  ERX13450982 |
| 4278  Large White | 03/06/2021 | SAMEA112465604 | 1000  Large White | normal breeding procedure | D9 | 18 | embryo_D9_8  embryo_D9_omic_pool_2 | SAMEA112465645  SAMEA115811555 | ovoid blastocyst 5mm | embryo_multiome_scRNAseq_D9_2  embryo_multiome_scATACseq_D9_2 | ERX13271548  ERX13271557  ERX13450983 |
| 4300  Large White | 03/06/2021 | SAMEA112465605 | 1000  Large White | normal breeding procedure | D9 | 3 | embryo_D9_9  embryo_D9_omic_pool_2 | SAMEA112465646  SAMEA115811555 | ovoid blastocyst 5mm | embryo_multiome_scRNAseq_D9_2  embryo_multiome_scATACseq_D9_2 | ERX13271548  ERX13271557  ERX13450983 |
| 4221  Large White | 03/06/2021 | SAMEA112465606 | 1000  Large White | normal breeding procedure | D11 | 1 | embryo_D11_4  embryo_D11_omic_pool_1 | SAMEA112465648  SAMEA115811556 | ovoid blastocyst 10-15mm | embryo_multiome_scRNAseq_D11_1  embryo_multiome_scATACseq_D11_1 | ERX13271549  ERX13271558  ERX13450984 |
| 4086  Large White | 03/06/2021 | SAMEA112465607 | 1000  Large White | normal breeding procedure | D11 | 1 | embryo_D11_6  embryo_D11_omic_pool_1 | SAMEA112465650  SAMEA115811556 | ovoid blastocyst 10-15mm | embryo_multiome_scRNAseq_D11_1  embryo_multiome_scATACseq_D11_1 | ERX13271549  ERX13271558  ERX13450984 |
| 4221  Large White | 03/06/2021 | SAMEA112465606 | 1000  Large White | normal breeding procedure | D11 | 6 | embryo_D11_5  embryo_D11_omic_pool_2 | SAMEA112465649  SAMEA115811557 | ovoid blastocyst 10-15mm | embryo_multiome_scRNAseq_D11_2  embryo_multiome_scATACseq_D11_2 | ERX13271550  ERX13271559  ERX13450985 |
| 4086  Large White | 03/06/2021 | SAMEA112465607 | 1000  Large White | normal breeding procedure | D11 | 6 | embryo_D11_7  embryo_D11_omic_pool_2 | SAMEA112465651  SAMEA115811557 | ovoid blastocyst 10-15mm | embryo_multiome_scRNAseq_D11_2  embryo_multiome_scATACseq_D11_2 | ERX13271550  ERX13271559  ERX13450985 |
| 2548  Large White | 02/02/2021 | SAMEA112465595 | 2000966  crossed | normal breeding procedure | D11 | 5 | embryo_D11_2  embryo_single_cell_D11_3 | SAMEA112465627  SAMEA115811551 | ovoid blastocyst 10-15mm | embryo_multiome_scRNAseq_D11_3  embryo_multiome_scATACseq_D11_3 | ERX13271551  ERX13271560  ERX13450986 |
| 2548  Large White | 02/02/2021 | SAMEA112465595 | 2000966  crossed | normal breeding procedure | D11 | 5 | embryo_D11_3  embryo_single_cell_D11_4 | SAMEA112465628  SAMEA115811552 | ovoid blastocyst 10-15mm | embryo_multiome_scRNAseq_D11_4  embryo_multiome_scATACseq_D11_4 | ERX13271552  ERX13271561  ERX13450987 |
