## Supplementary Table 2 for "Single-cell omics reveal distinct gene regulatory dynamics underpinning embryonic and extraembryonic lineage functions during pig blastocyst development"

| **Samples** | **Number of raw RNA-seq reads** | **% of RNA-seq reads confidently mapped to the genome** | **Median UMIs counts per Cell** | **Number of detected genes** | **Median high-quality fragment per cell** | **Number of raw ATAC-seq reads** | **TSS enrichment score** | **% of ATAC-seq reads confidently mapped to the genome** | **Number of cells before quality controls** | **Number of cells after quality controls** |
| --- | --- | --- | --- | --- | --- | --- | --- | --- | --- | --- |
| E7 | 245M | 92.1 | 1,800 | 28,601 | 12,656 | 433M | 2.54 | 87.3 | 2,426 | 894 |
| E9-1 | 263M | 83.8 | 9,430 | 28,935 | 2,840 | 118M | 2.37 | 87.1 | 3,074 | 1,544 |
| E9-2 | 267M | 83.7 | 7,601 | 28,987 | 5,324 | 304M | 1.92 | 86.9 | 3,032 | 1,673 |
| E11-1 | 332M | 76.9 | 9,330 | 28,705 | 7,848 | 197M | 3.08 | 86.2 | 3,524 | 2,338 |
| E11-2 | 407M | 75.3 | 9,420 | 29,815 | 5,396 | 207M | 2.73 | 85.9 | 4,468 | 2,807 |
| E11-3 | 283M | 79.1 | 3,930 | 26,025 | 2,238 | 129M | 2.18 | 85.6 | 2,353 | 1,023 |
| E11-4 | 271M | 79.8 | 3,649 | 28,260 | 6,292 | 308M | 2.34 | 86.3 | 4,679 | 2,599 |
